# Reconstructing tumor evolution without chromosomal instability reveals in vivo determinants of therapeutic response

**DOI:** 10.1101/2025.03.17.643814

**Authors:** Ruhollah Moussavi-Baygi, Matthew J. Ryan, Woogwang Sim, Samuel B. Hoelscher, Valbona Luga, Arun A. Chandrakumar, Lore Hoes, Junghwa Cha, Young Sun Lee, Katelyn Herm, Ben Doron, Danika Bakke, Sharon Wu, C.K. Cornelia Ding, Bradley A. Stohr, Peng Jin, Tejasveeta Nadkarni, Xiangyi Fang, Melita Haryono, An Nguyen, Wouter R. Karthaus, Charles L. Sawyers, Felix Y. Feng, Hani Goodarzi, Rohit Bose

## Abstract

Tumor responses to therapy are profoundly shaped by selective pressures operating in vivo, including tumor-host interactions, tissue architecture, and endocrine signaling, yet most genome-scale approaches are conducted in vitro. We developed Stochastically Emergent Tumors (SETs), an in vivo platform that distills tumor evolution into sparse genetic variation amenable to computational inference and deep learning while uniquely avoiding chromosomal instability. Unlike single-perturbation screens, SETs evolve through combinatorial alterations, including neomorphic variants, across the entire genome. Applied to prostate cancer, SETs revealed how hormone therapy reshapes tumor evolution, identifying six unrecognized sensitization determinants, including ZFHX3 loss, together with resistance-associated alterations in CIC and KMT2D. Patient tumor analyses linked these potential biomarkers to clinical outcomes and nominated ZFHX3 as a target to push tumors into a luminal state. By decoupling tumor evolution from chromosomal instability, SETs harness in vivo therapeutic selection as a genome-scale discovery engine for determinants of both sensitivity and resistance.

**Statement of Significance:** Most genome-scale cancer screens cannot capture in vivo therapeutic selection. Stochastically Emergent Tumors enable evolution without chromosomal instability, generating sparse genetic variation amenable to deep learning. This transforms tumor evolution into a genome-scale discovery engine for determinants of therapeutic sensitivity, exemplified by ZFHX3 loss promoting hormone-responsive prostate cancer.

## Introduction

Cancer therapy response is determined not only by effects on cellular proliferation but also by selective pressures imposed by the tumor microenvironment, including hormonal, immune, and architectural cues. Genome-scale in vitro interrogation of transformed cancer cells has identified numerous correlates of therapeutic response, but these predominantly capture cell-autonomous mechanisms of resistance. By contrast, determinants of therapeutic sensitivity linked to differentiation state and microenvironmental context remain poorly resolved at genome scale. This limitation is particularly evident in hormone-dependent cancers, where therapeutic response is tightly coupled to epithelial lineage identity, including luminal and basal differentiation states that are difficult to interrogate at genome scale in vitro. As a result, a substantial class of clinically relevant determinants governing therapeutic sensitization and lineage-dependent vulnerability remains systematically underexplored.

Interrogation of therapeutic evolution is fundamentally limited by the genomic complexity of tumors. Genome-scale experiments typically rely on predefined knockout libraries to interrogate single-gene loss-of-function in already transformed cancer cells. These approaches are difficult to scale in vivo and poorly suited to interrogate allelic diversity and non-coding variation under physiological selection. In vitro screens, while experimentally tractable, incompletely model the selective pressures imposed by the tumor microenvironment and predominantly select for resistant phenotypes^1^. In parallel, analyses of patient tumors aim to identify enriched or depleted alterations following therapy but are confounded by co-alteration of neighboring genes due to chromosomal instability^2,3^. Together, these approaches interrogate therapeutic evolution in tumors shaped by chromosomal instability, which obscures inference of the coding and non-coding variants underlying therapeutic response.

These limitations raise a fundamental question: if chromosomal instability were constrained, could tumor evolution itself serve as a genome-scale mechanism for discovering determinants of therapeutic response? Rather than imposing predefined perturbations, physiological selective pressures within the tumor microenvironment would enable genome-wide exploration of genetic variation through mutation and selection. Physiological selection would encode therapeutic fitness directly into sparse genetic variants amenable to computational inference. Prior approaches to accelerate mutagenesis, including carcinogen exposure or APOBEC activation, also induce chromosomal instability, thereby obscuring the evolutionary information required to infer specific variants driving therapeutic response.

Here, we establish a system that reconstructs tumor evolution without chromosomal instability. By initiating tumorigenesis from genetically defined non-malignant organoid substrates and enabling evolution through sparse point mutations, complex selective pressures imposed by the tumor microenvironment are distilled into interpretable genomic variation. Because this system is not constrained by predefined knockout libraries, it enables genome-wide interrogation of coding and non-coding variants while permitting combinatorial evolution under physiological selection. The resulting sparse mutational landscape is directly amenable to computational inference, including deep learning–based prediction of variant function. Applied to prostate cancer, this approach reveals in vivo determinants of therapeutic response, including previously obscured drivers of lineage-associated sensitivity and resistance.

### Mismatch repair deficiency reveals high-resolution evolutionary information obscured by chromosomal instability

Inferring determinants of therapeutic evolution remains challenging, even in large cross-sectional patient cohorts. Tumor genomes are often dominated by chromosomal instability, resulting in copy number alterations that span multiple genes^4^ (**Figure 1a**). For example, genomic analysis of prostate cancer patients treated with mainstay Androgen Receptor Pathway Inhibitors (ARPI) revealed enrichment at Xq12 encompassing the target Androgen Receptor (*AR*) gene^5–7^ (**Figures 1a-b, Supplementary Fig. 1a-b**). Beyond this locus, few additional alterations distinguished ARPI-treated from ARPI-naïve tumors. Notably, >95% of these copy number alterations were multigenic (**Supplementary Fig. 1c**). This multigenic structure obscures inference of individual gene contributions within co-altered regions, such as loss of *ZFHX3* versus *PHLPP2* (16q22) or loss of *CIC* versus *ERF* (19q13)^8–11^ (**Figure 1a**). In contrast, tumor subgroups with low chromosomal instability acquired discrete point mutations within genes that are typically co-altered by copy number changes (**Figure 1c**, **Supplementary Fig. 1d**). This enables higher-resolution inference within otherwise co-altered loci: focusing on the low FGA^12–14^ tumors reveals ASXL1 and DNMT3B as the only genes harboring putative driver mutations within the 11-gene amplicon at 20q11 (**Figure 1c**). Accordingly, tumor cohorts that evolve predominantly through point mutations in the absence of chromosomal instability permit high-resolution inference of tumor evolution.

**Figure 1.**
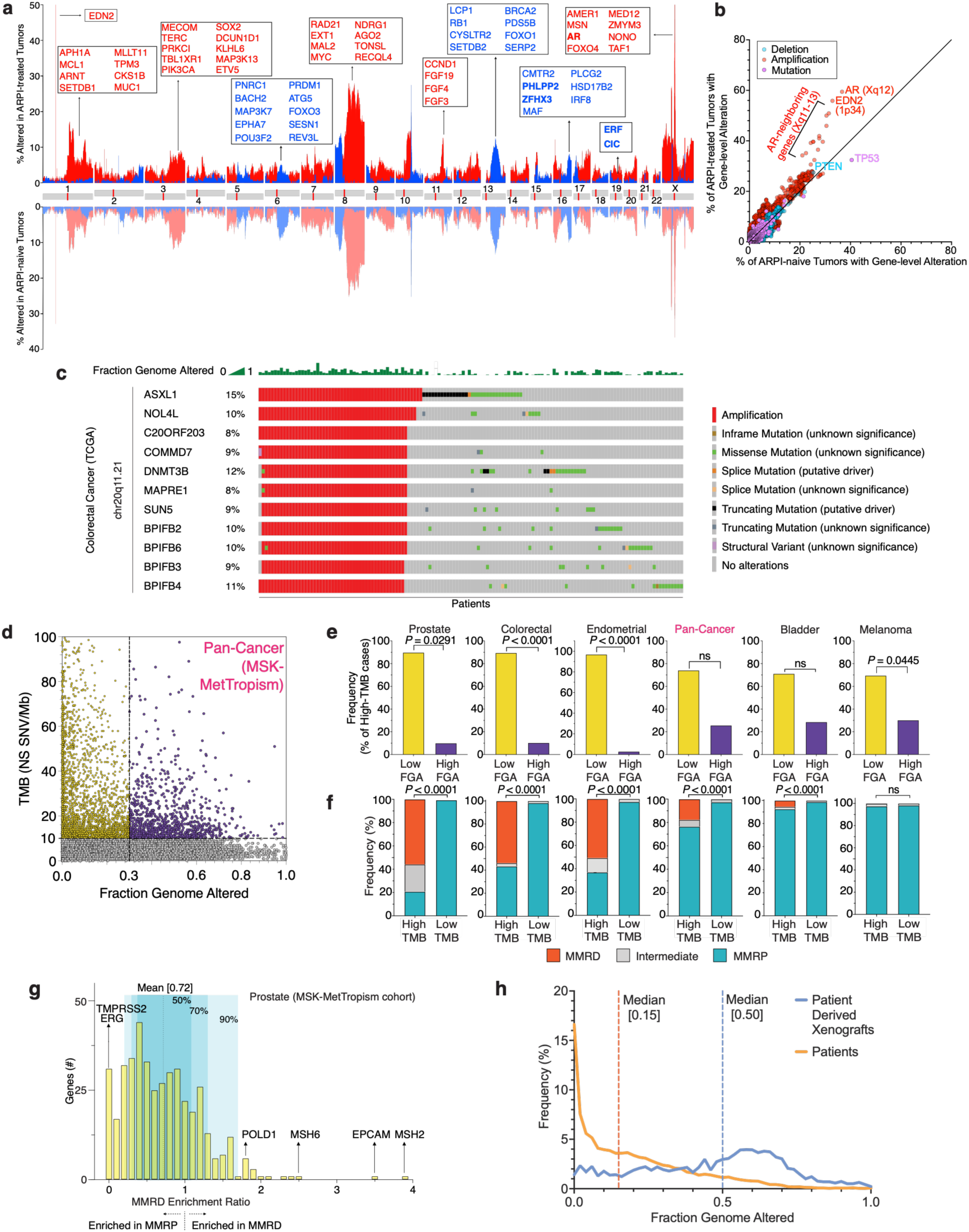
– Mismatch repair deficiency enables high-resolution inference of tumor evolution obscured by chromosomal instability. (**a**) Landscape of copy number alterations in ARPI-treated (*top*) and ARPI-naïve (*bottom*) advanced prostate cancer patient tumors. The frequency of copy number alterations is graphed versus chromosomal locus. Copy number alterations are amplifications (red) and deep deletions (blue)^2^. Examples of OncoKB annotated cancer-related genes^137,138^ are highlighted at loci subject to frequent copy number alterations. Whole exome sequencing (WES) results are from the PCF/SU2C East Coast Dream Team metastatic castration-resistant prostate cancer cohort^5^ (*n* = 444). (**b**) The rate of deletion (blue), amplification (red), and mutation (magenta) of genes is compared in ARPI-treated tumors versus ARPI-naïve tumors in the PCF/SU2C East Coast Dream Team metastatic castration-resistant prostate cancer cohort^5^ (*n* = 444). (**c**) Point mutation pattern in cancer patients with high FGA tumors (mostly on the left) compared to low FGA tumors. The OncoPrint is derived from colorectal cancer patients (*n* = 594) in the TCGA cohort WES^128^ obtained from cBioPortal^134–136^. **(d)** Tumor Mutation Burden (TMB) is graphed versus Fraction of Genome Altered (FGA) by copy number alterations, for each patient’s tumor, across cancer types. The dotted lines indicate high-TMB (10 NS SNVs / Mb) and high-FGA (0.3) thresholds. Targeted panel sequencing from MSK-MetTropism cohort^33^: endometrial (*n =* 366), prostate (*n =* 79), colorectal (*n =* 595), small bowel (*n =* 18), non-melanoma skin (*n =* 79), hepatobiliary (*n =* 44), cervical (*n =* 25), anal (*n =* 16), head and neck (*n =* 74), pan-cancer (*n =* 24,753), pancreatic (*n =* 39), bladder (*n =* 459), melanoma (*n =* 543), esophagastric (*n =* 55), non-small cell lung (*n =* 1115), breast (*n =* 129), ovarian (*n =* 26), and small cell lung (*n =* 108). **(e)** Cancer-specific frequency of high-TMB tumors that have low FGA (≤0.3) versus high FGA. Statistical significance by one-sided hypergeometric test. Data from MSK-MetTropism cohort described above. **(f)** Cancer-specific frequency of mismatch repair deficiency (MMRD) or mismatch repair proficiency (MMRP) among High-TMB (≥10 NS SNVs/Mb) and Low-TMB tumors. MMR status is categorized using MSIsensor score^124^ and color-coded: red (≥10, MMRD), grey (between 4-10, intermediate), or blue (≤4, MMRP). A one-sided hypergeometric test was performed to compare the percentage of MMRD tumors between groups. Data from MSK-MetTropism cohort described above. Additional analysis in **Supplementary Figure 1e**. **(g)** Relative frequency of gene variants compared between MMRD and MMRP patient tumors. For each gene, an MMRD Enrichment Ratio is calculated as TMB-normalized frequency of variants (point mutations, copy number alterations, structural variants) in MMRD versus MMRP tumors, details in **Methods**. A histogram of the MMRD Enrichment Ratio is graphed for all genes, with 50%, 70%, and 90% of all genes shown centered around the mean value (0.72). The bin size is 0.1. Individual genes are highlighted within the most MMRD and MMRP enriched bins. Targeted panel sequencing from MSK-MetTropism^33^ was analyzed with MMRP (*n =* 1,831) and MMRD (*n =* 43) prostate cancer patients. **(h)** Histogram of FGA distribution, comparing pan-cancer patient tumors with patient-derived xenografts (PDXs). Data from patients (n = 24,753) and cell lines (n = 6,272) from MSK-MetTropism and NCI Patient-Derived Models Repository cohort, respectively. The histograms of FGA are graphed with a bin size of 0.02. The median FGAs in patients (0.15) and PDXs (0.50) are highlighted. Statistical significance by two-tailed, unpaired Welch’s t-test and two-tailed, unpaired Mann-Whitney test.

To identify patient tumor evolutionary states that maintain low chromosomal instability, we examined the relationship between point mutations and FGA across cancer types (**Figure 1d**). A high Tumor Mutation Burden (TMB) ≥10 was significantly associated with a low FGA (≤0.3) among prostate, endometrial, and colorectal cancers^15,16^ (**Figure 1e**). These cancer types in which high TMB is predominantly driven by DNA mismatch repair deficiency (MMRD)^17^ (**Figure 1f, Supplementary Fig.1e-f**). By contrast, high TMB tumors driven by non-MMRD mutagenesis, including bladder cancer and melanoma, exhibited higher FGA^18–22^ and retained recurrent multigenic chromosomal alterations (**Figure 1e-f, Supplementary Fig. 2**). Given the uniquely low FGA associated with MMRD, we focused on MMRD tumor evolution. We directly compared MMRD to mismatch repair proficient (MMRP) tumor evolution and found that most genes had similar relative alteration frequencies within each cancer type, irrespective of mutational mechanism^23,24^ (**Figure 1g**, **Supplementary Fig. 1h**). As expected, MMRD-enriched exceptions included *MSH2* and *MSH6*, the loss of both of which initiates MMRD^16,25,26^ whereas MMRP-enriched exceptions included genes primarily affected by structural variants rather than point mutations^27,28^ (**Figure 1g, Supplementary Fig. 1g**). Collectively, these findings indicate that MMRD mutagenesis enables tumors to evolve through alterations in similar genes, but without chromosomal instability, thereby revealing evolutionary information otherwise obscured by multigenic copy number alterations. Because xenograft tumors exhibit even greater chromosomal instability than patient tumors (**Figure 1h**), this motivated the development of a tumor evolution system that avoids chromosomal instability.

### Stochastically emergent tumors generate heterogeneous evolutionary trajectories absent from conventional models

To reconstruct malignant evolution without inducing chromosomal instability or relying on extrinsic carcinogens, we tested whether mismatch repair deficiency alone could drive tumorigenesis from non-malignant organoid substrates^29–32^. Based on features of mismatch repair–deficient (MMRD) patient prostate tumors^22^ (**Supplementary Fig. 1g**), we used CRISPR to knock out *Msh2* in a murine prostate organoid line to establish MMRD lines, which we termed “pregraft” lines (**Figure 2a**). To accelerate tumorigenesis, we also generated Pten^KO^;Msh2^KO^ and Trp53^KO^;Msh2^KO^ pregraft lines, reflecting two of the most frequently altered tumor suppressors in prostate cancer^33^ (**Supplementary Fig. 3a-c**). Prior studies have shown that Pten^KO^ or Trp53^KO^ alone are insufficient to yield malignancy^9,34–39^. MMRD pregraft lines exhibited high intra-sample mutational heterogeneity (**Supplementary Fig. 3d-e**) and were injected subcutaneously into SCID mice^9^. The resulting MMRD tumors maintained patient-like TMBs and low FGAs, thereby preserving interpretable tumor evolution without chromosomal instability (**Figure 2b-c**). Notably, these tumors maintained substantially sparser mutational landscapes than commonly used AR-dependent prostate cancer cell lines (LNCaP, 22RV1, MDA-PC-2b) ^31,32^ (**Figure 2c**). By contrast, none of the MMRP Pten^KO^ or Trp53^KO^ negative control pregrafts formed tumors. Because the MMRD tumors emerged without extrinsic carcinogens or engineered oncogenic combinations, we refer to them as Stochastically Emergent Tumors (SETs).

**Figure 2.**
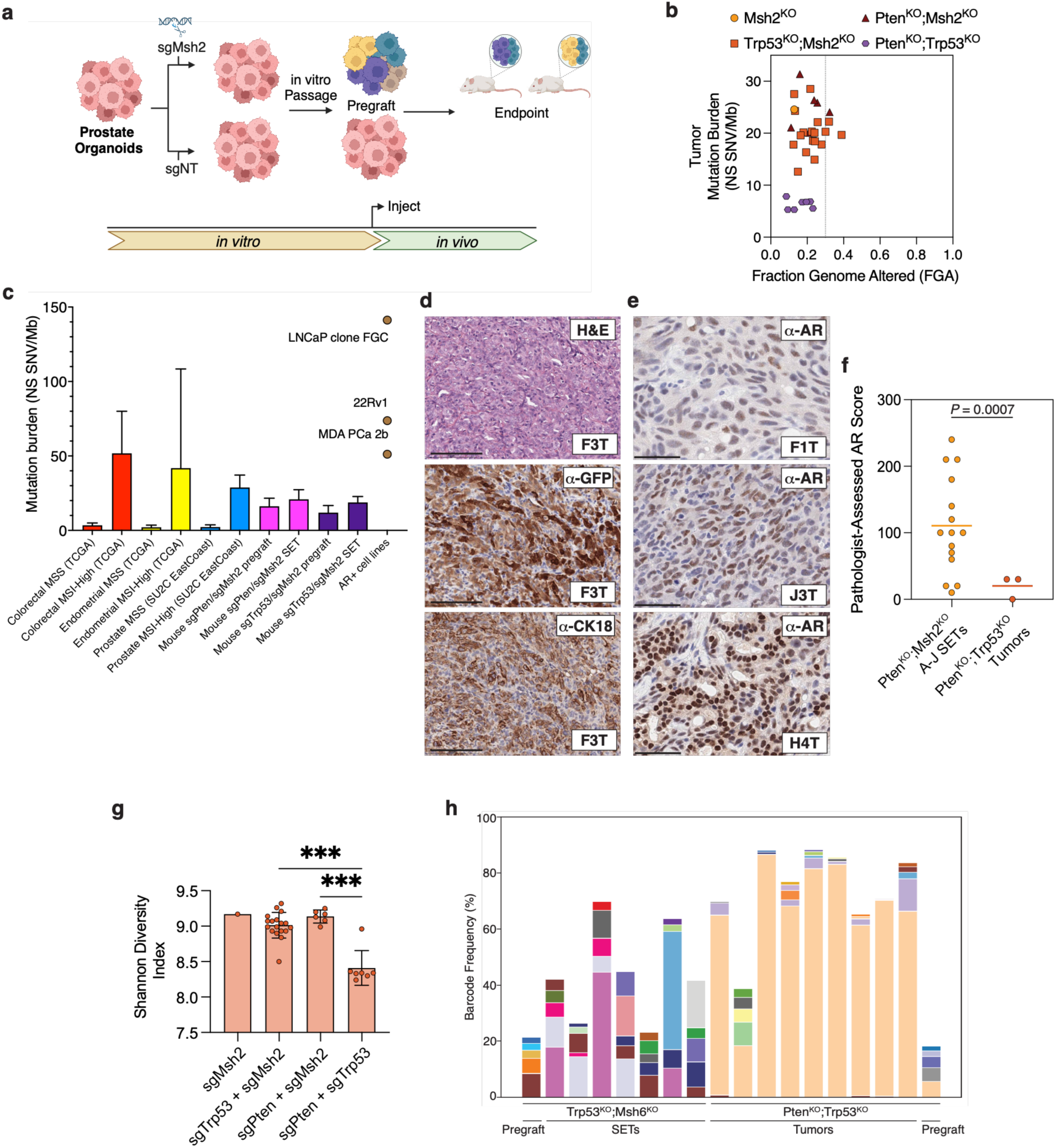
– Stochastically Emergent Tumors (SETs) generate heterogeneous evolutionary trajectories without chromosomal instability. (**a**) Experimental scheme to determine whether MMRD organoids become tumorigenic. A prostate organoid line was subjected to CRISPR/Cas9 knockout with various guides, including sgMsh2 or a non-targeting guide (sgNT). The sgMsh2-nucleofected (*Msh2*^KO^) organoids were passaged in vitro for 355 days, then injected subcutaneously into mice, which were observed for 8 additional months. (**b**) Analysis of mutation burden versus fraction genome altered for tumors resulting from murine injection of MMRD or MMRP pregrafts. Resulting tumors analyzed: *Msh2*^KO^-alone, *n =* 1; *Trp53*^KO^*;Msh2*^KO^, *n =* 18; *Pten*^KO^*;Msh2*^KO^, *n* = 6; *Pten*^KO^*;Trp53*^KO^, *n =* 7. (**c**) Comparison of the mutation burdens of patient tumors, SETs, pregrafts, and 3 AR expressing prostate cancer cell lines. Median, interquartile range, and whiskers graphed per Tukey’s method. Patient data includes patients with TMB from 10 to 50 NS SNV/Mb and MSISensor score > 10 for colorectal (*n* = 65) and endometrial cancer (*n* = 122, TCGA^128^), and patients with deletion or mutation of *MSH2* or *MSH6* in the prostate (*n* = 8, SU2C EastCoast^5^). (**d**) Representative SET (F3T) derived from *Pten*^KO^*;Msh2*^KO^ subline F was subjected to hematoxylin and eosin (H&E) staining, as well as immunohistochemistry (IHC) for Green Fluorescent Protein (GFP) from the organoid pregraft line and the luminal epithelial marker cytokeratin 18 (CK18). Scale bars equal 100 mm. Additional examples in **Supplementary Figure 4d**. (**e**) Nuclear hormone receptor levels among SETs arising from different *Pten*^KO^*;Msh2*^KO^ MMRD clones. Representative immunohistochemistry (IHC) for the androgen receptor (AR) of 3 distinct SETs (F1T, H4T, J3T) arising from *Pten*^KO^*;Msh2*^KO^ F, H, and J subline injections. Scale bars equal 50 mm. Additional IHC in **Supplementary Figure 4f**. (**f**) Pathologist quantification of nuclear AR among SETs arising from different MMRD clones. A genitourinary pathologist assessed an AR score for each SET, as well as control MMRP *Pten*^KO^*;Trp53*^KO^ tumors based on the nuclear AR intensity (0-3+) multiplied by the percentage of AR positive cells. P-value by Welch’s unpaired t-test. (**g**) Shannon Diversity Index for *Msh2*^KO^-alone, *n =* 1; *Trp53*^KO^*;Msh2*^KO^, *n =* 18; *Pten*^KO^*;Msh2*^KO^, *n* = 6; and *Pten*^KO^*;Trp53*^KO^, *n =* 7 tumors. (**h**) Comparison of the barcoded clonal composition of SETs versus control MMRP tumors, as well as their cognate pregrafts. Frequency of the top 5 barcoded lineages within each sample, with unique colors for distinct barcodes. Samples analyzed by barcode sequencing: *Trp53*^KO^*;Msh6*^KO^ (sgTrp53+sgMsh6) SETs *n* = 7; cognate *Trp53*^KO^*;Msh6*^KO^ pregraft *n* = 1, MMRP *Pten*^KO^*;Trp53*^KO^ (sgPten+sgTrp53) tumors *n* = 9; cognate *Pten*^KO^*;Trp53*^KO^ pregraft *n* = 1.

High intra- and inter-tumoral heterogeneity generated through tumor evolution underlies variable therapeutic responses in human cancers^40^. By contrast, tumors derived from conventional xenograft and genetically engineered mouse models exhibit constrained evolutionary heterogeneity^41,42^. To assess heterogeneity in SETs, we generated multiple independent Pten^KO^;Msh2^KO^ pregraft sublines derived from a common parental population (**Supplementary Fig. 4a-b, Supplementary Table 1**). Several sublines reproducibly formed tumors with short latencies, enabling practical timelines for in vivo studies (**Supplementary Fig. 4c**). SETs retained luminal epithelial marker expression yet exhibited pronounced inter- and intra-tumoral heterogeneity in nuclear AR expression^30,39,43–46^ (**Figure 2d-e, Supplementary Fig. 4d-f**). SETs also displayed significantly higher nuclear AR levels than aggressive-variant Pten^KO^;Trp53^KO^ tumors^47^ (**Figure 2e-f, Supplementary Fig. 4f**). Importantly, MMRD and MMRP pregrafts exhibited similar, uniform AR protein levels prior to implantation, indicating that AR heterogeneity emerged during in vivo tumor evolution (**Supplementary Fig. 4e**). Thus, stochastic mutational heterogeneity within MMRD pregrafts (**Supplementary Fig. 3e**) generates SETs with marked phenotypic heterogeneity in growth kinetics and endogenous therapeutic target expression.

As expected, SETs demonstrated significantly higher mutational heterogeneity than conventional tumor models (**Figure 2g**). To directly compare the clonal architectures underlying this evolutionary heterogeneity, we used DNA barcode-based lineage tracing^48^. Trp53^KO^;Msh6^KO^ SETs were generated to complement our analysis of Pten^KO^;Msh2^KO^ SETs (**Supplementary Fig. 5a**) and exhibited high intra- and inter-tumoral clonal heterogeneity by lineage tracing (**Figure 2h, Supplementary Fig. 5b-c**). By contrast, MMRP Pten^KO^;Trp53^KO^ control tumors showed markedly reduced clonal diversity (**Figure 2h**), with a single dominant lineage comprising a median 68% of each tumor^49–51^. Dominant clonal lineages in SETs were distinct from those present in their corresponding pregraft, indicating de novo clonal selection in vivo (**Figure 2h**). Together, these findings demonstrate that SETs undergo stochastic endogenous clonal evolution, generating genetic and phenotypic heterogeneity in sharp contrast to the clonal dominance characteristic of conventional tumor models.

### SETs reveal genome-wide determinants of in vivo hormone therapy response, exemplified by Zfhx3 loss

Because SETs evolve in vivo without chromosomal instability, we asked whether their mutational and clonal trajectories would reconfigure under therapeutic pressure, revealing determinants of divergent therapeutic response. To model first-line hormone therapy, we treated SET-bearing mice with surgical or chemical castration, with or without androgen receptor pathway inhibition (ARPI)^52–58^. Pten^KO^;Msh2^KO^ or Trp53^KO^;Msh2^KO^ SETs evolved under androgen-intact conditions, castration, or castration plus ARPI^9,59,60^ (**Figure 3a**). Pten^KO^;Msh2^KO^ SETs exhibited significantly greater in vivo responses to hormone therapy than Trp53^KO^;Msh2^KO^ SETs (**Figure 3b, Supplementary Fig. 6a-b**), consistent with clinical observations^5,61–64^. SET-specific mutations clustered remarkably by both truncal genotype and hormone therapy status (**Supplementary Fig. 6c**). In the absence of lineage barcoding, clonal structure was inferred by unsupervised phylogenetic analysis of SET-specific mutations^65–69^ (**Figure 3c**). Notably, Trp53^KO^;Msh2^KO^ SETs developed distinct mutational and clonal identities under castration despite minimal macroscopic tumor response (**Figure 3c, Supplementary Fig. 6b-c**). To directly test therapy-driven evolutionary divergence, barcoded Trp53^KO^;Msh6^KO^ SETs were treated with castration or vehicle control. Castration-resistant SETs exhibited distinct clonal dynamics and significantly increased inter-tumoral heterogeneity compared with androgen-intact SETs (**Figure 3c-d**). Together, these findings demonstrate that SETs evolve divergently under therapeutic selection in a genotype-dependent manner, even in the absence of macroscopic tumor response.

**Figure 3.**
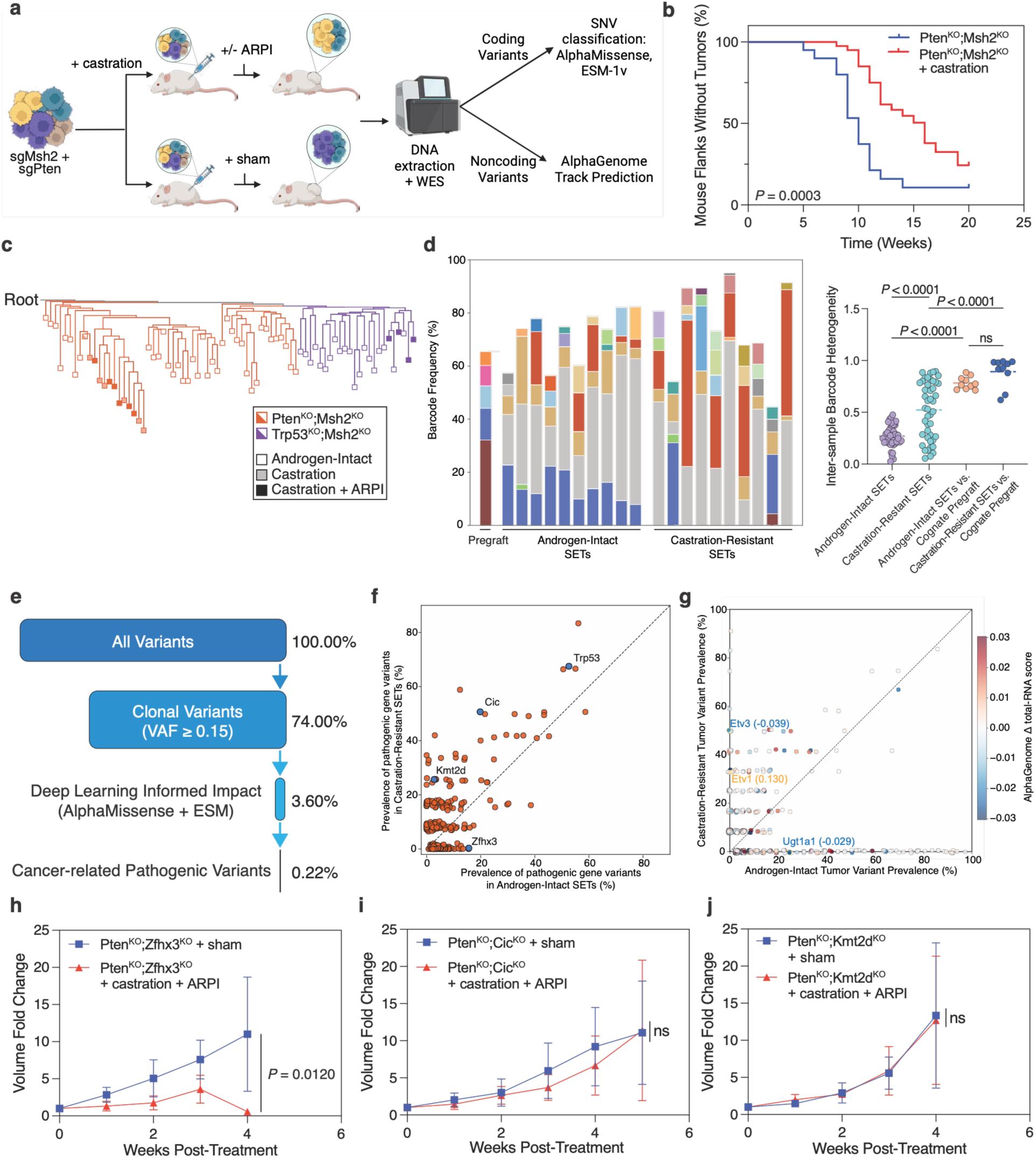
– SETs reveal genome-wide determinants of in vivo hormone therapy response. (**a**) Experimental scheme for genome-wide interrogation of coding and non-coding variants associated with in vivo hormone therapy response. *Pten*^KO^;*Msh2*^KO^ SETs were subjected to castration with or without AR pathway inhibition (ARPI), followed by whole-exome sequencing and computational prioritization of coding and non-coding variants using AlphaMissense, ESM-1v, and AlphaGenome. **(b)** Effect of castration on SET growth kinetics. Kaplan-Meier of mouse flanks without tumors ≥ 200 mm^3^ for *Pten*^KO^;*Msh2*^KO^ (sgPten+sgMsh2) SETs upon castration. Androgen-intact mice *n =* 10; castrated mice *n =* 20. P-value by log-rank test. (**c**) Relationship of SET clones upon in vivo cancer therapy and with distinct truncal genotypes. Phylogenetic tree of inferred clones from *Pten*^KO^;*Msh2*^KO^ genotype: pregrafts (*n =* 21); androgen-intact SETs (*n* = 36); castration-alone SETs (*n* = 6); and castration + ARPI SETs (*n* = 6); *Trp53*^KO^;*Msh2*^KO^ genotype: pregrafts (*n* = 15); androgen-intact SETs (*n* = 21); castration + ARPI SETs (*n* = 4). (**d**) Evaluation of inter-tumoral heterogeneity among lineage-traced castration-resistant SETs compared to androgen-intact SETs. Inter-sample mutational heterogeneity was assessed by Bray-Curtis Dissimilarity of barcode lineage composition^139^. Plot shows indicated pairwise comparisons within the *Trp53*^KO^*;Msh6*^KO^ androgen-intact (*n* = 10) and castration-resistant (*n* = 10) SET groups, along with comparisons to the cognate pregraft (*n* = 1). P-values by Welch’s one-way ANOVA with post hoc Dunnett’s T3 multiple comparisons test (ns = not significant). (**e**) Identification of candidate drivers of divergent therapeutic evolution in SETs. Variants were filtered sequentially based on clonality (VAF ≥ 0.15), predicted functional impact by AlphaMissense or ESM-1v/2, and presence within cancer-related genes on clinically relevant targeted sequencing panels^140–142^. (**f**) Comparison of pathogenic gene variant prevalence between castration-resistant SETs versus androgen-intact *Pten*^KO^*;Msh2*^KO^ SETs. For each putative driver, the prevalence in castration-resistant SETs (*n* = 12) containing a point mutation is graphed versus the corresponding prevalence in androgen-intact SETs (*n* = 36), with jitter for visualization. (**g**) Comparison of noncoding mutations related to genes in clinically targeted cancer panels between castration-resistant versus androgen-intact *Pten*^KO^*;Msh2*^KO^ SETs. For each mutation, the prevalence in castration-resistant SETs (*n* = 12) is graphed versus androgen-intact SETs (*n* = 36), with jitter for visualization. Details in **Methods**. (**h**) Interrogation of hormone therapy response upon *Zfhx3* knockout. Tumor volume fold change upon castration + ARPI versus sham treatment is graphed following murine injection of prostate organoids with CRISPR knockout of *Pten*^KO^*;Zfhx3*^KO^ (sgPten+sgZfhx3). (*n =* 5 mice per arm). P-value by mixed-effects model demonstrating the significance of the interaction between treatment and weeks. (**i**) Interrogation of hormone therapy response upon *Cic* knockout. Same as **h** with CRISPR knockout of *Pten*^KO^*;Cic*^KO^ (sgPten+sgCic). ns, not significant by mixed-effects model demonstrating significance of the interaction between treatment and weeks. (**j**) Interrogation of hormone therapy response upon *Kmt2d* knockout. Same as **h** with CRISPR knockout of *Pten*^KO^*;Kmt2d*^KO^ (sgPten+sgKmt2d). ns, not significant by mixed-effects model demonstrating significance of the interaction between treatment and weeks.

Because SETs evolve through stochastic point mutations, they enable tractable whole-genome interrogation of genetic variants associated with in vivo therapeutic response (**Supplementary Fig. 6d**). Deep learning–informed filtering prioritized a small subset of candidate coding and non-coding pathogenic drivers from passenger mutations^70^ (**Figure 3e-g**). As a positive control, both androgen-intact and castration-resistant Pten^KO^;Msh2^KO^ SETs acquired recurrent *Trp53* pathogenic variants, consistent with its dual role as a canonical tumor suppressor and driver of hormone therapy resistance in prostate cancer. Several pathogenic variants were selectively enriched during androgen-intact tumor evolution, implicating candidate determinants of hormone sensitivity (**Figure 3e-f**). Notably, Pten^KO^;Msh2^KO^ androgen-intact SETs were enriched for loss-of-function mutations in *Zfhx3*, a zinc-finger homeobox tumor suppressor (**Figure 3f, Supplementary Fig. 7a**) that binds and represses androgen receptor signaling^8,11,71,72^. We next examined SET-specific mutations enriched during castration-resistant tumor evolution. Castration-resistant SETs were enriched for loss-of-function mutations in *Cic* and *Kmt2d*, genes implicated in lineage regulation and plasticity across cancer types (**Figure 3e-f**). Consistent with this, MSI-high prostate tumors similarly exhibited focal *CIC* mutations independent of *ERF* (**Supplementary Fig. 9b**). Extending beyond coding regions, computational analysis of recurrent non-coding variants with AlphaGenome identified predicted transcriptional regulatory alterations linked to divergent therapeutic evolution, including pathogenic variants associated with repression of the androgen-glucuronidating enzyme *UGT1A1* in androgen-intact SETs, consistent with the retention of intracellular androgen (**Figure 3g**). Similarly, AlphaGenome predicted activation of oncogenic *ETV1* alongside suppression of repressive *ETV3* in castration-resistant SETs. Moreover, these predictions were consistent with corresponding transcriptional changes observed in SET transcriptomes^38,73–77^ (**Supplementary Fig. 6e**). Collectively, these analyses demonstrate that SETs enable genome-wide in vivo identification of coding and noncoding pathogenic variants, distinguishing hormone-sensitive from castration-resistant tumor evolution.

To test whether SET-identified genes directly modulate hormone sensitivity, we engineered Pten^KO^;Zfhx3^KO^ organoids and compared them to Pten^KO^;Kmt2d^KO^ and Pten^KO^;Cic^KO^ controls (**Supplementary Fig. 8a-c**). All three genotypes drove malignant evolution from non-malignant substrates, extending prior observations in transformed cell models to demonstrate bona fide tumorigenic capacity^8,11,71,72^. Consistent with SET-based predictions, Pten^KO^;Zfhx3^KO^ tumors exhibited marked sensitivity to hormone therapy, whereas Pten^KO^;Cic^KO^ and Pten^KO^;Kmt2d^KO^ tumors were castration-resistant (**Figure 3h-j**). Collectively, these findings establish *Zfhx3* loss as a determinant of hormone sensitivity and demonstrate the capacity of SETs to identify genome-wide determinants of in vivo therapeutic response, including lineage-associated therapeutic states that have historically been difficult to interrogate at genome scale.

### SET-identified alterations such as ZFHX3 loss define luminal hormone-responsive states associated with improved survival in prostate cancer patients

We next asked whether *ZFHX3* loss-of-function alterations identified by SETs stratify clinical outcomes in prostate cancer patients, the vast majority of whom have mismatch repair–proficient (MMRP) tumors. We analyzed inferred overall survival in a large cohort of MMRP prostate adenocarcinoma patients profiled by Caris Life Sciences, of whom 74% were documented to have received hormone therapy. Patients with *ZFHX3*-altered tumors exhibited markedly longer inferred overall survival (median 110 months) compared with patients harboring alterations in *KMT2D* (61 months), *TP53* (42 months), or *CIC* (37 months) (**Figure 4a**). The favorable outcomes associated with *ZFHX3* alterations in MMRP patient tumors support the broader relevance of SET-identified determinants beyond MMRD, consistent with our earlier observation that MMRD and MMRP tumors converge on similar gene-level evolutionary trajectories (**Figure 1g**). Few genomic alterations have been robustly associated with favorable clinical outcomes in prostate cancer: *SPOP* mutations are associated with more indolent disease, and *TMPRSS2–ERG* fusions mark a luminal transcriptional program^78–80^. We discovered that patients with *ZFHX3*-altered tumors exhibit comparably favorable inferred overall survival (**Figure 4b**). Multivariate analysis confirmed that *ZFHX3* and *SPOP* alterations were independently associated with longer inferred overall survival (**Supplementary Fig. 9c**). Notably, *ZFHX3*, *KMT2D*, and *CIC* alterations were more frequent among patients of African and Chinese ancestry than among patients of European ancestry, which may have limited their detection in prior analyses (**Figure 4c and Supplementary Fig. 9a**).

**Figure 4.**
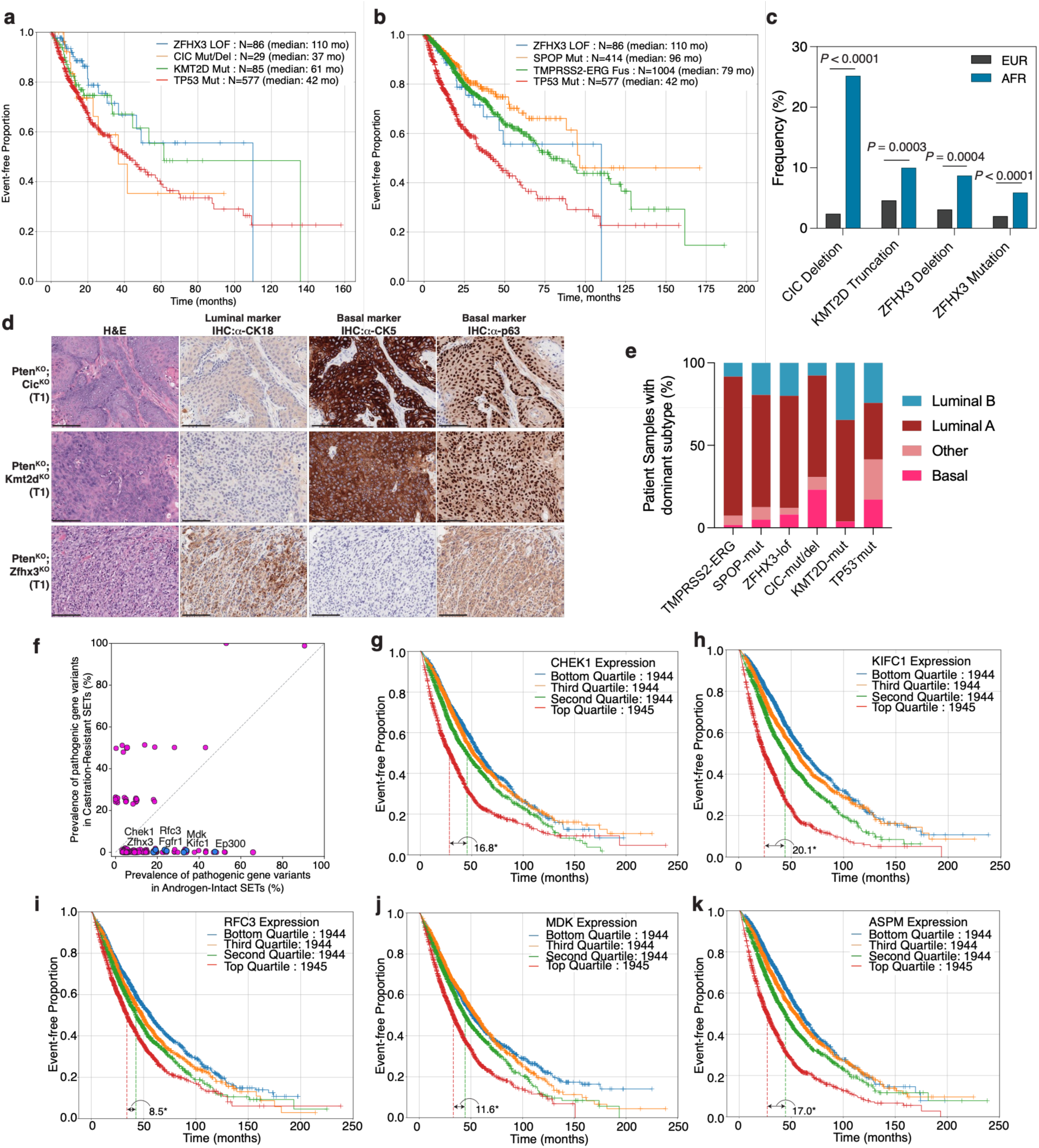
– SET-identified alterations define lineage-associated hormone-response states linked to prostate cancer survival. (**a**) and **(b)** Inferred overall survival in prostate cancer patients with the indicated genotypes. From a multicenter cohort of patients with prostate adenocarcinoma profiled by Caris Life Sciences. LOF – loss-of-function, Mut – mutated, Del – deleted. Performance characteristics in **Supplementary Table 2**. (**c**) Comparison of gene variant frequency between prostate cancer patients who are African American (AFR) versus those in a predominantly European American (EUR) cohort. The rate of *CIC* deep deletions was compared in AAPC^143^ and TCGA^128^ cohorts. The relative frequencies of *ZFHX3* and *KMT2D* variants were reported from the above and the Foundation combined primary and metastatic prostate cancer cohort^76^. Cohort sizes: AAPC *n* = 99, TCGA *n* = 494, Foundation AFR *n* = 288, Foundation EUR *n* = 2303. P-values by binomial test. (**d**) Luminal and basal marker analysis of tumors with differential response to hormone therapy. Histology and immunohistochemistry (IHC) of representative androgen-intact tumors (T1) arising from murine injection of prostate organoids with CRISPR knockout genotypes: *Pten*^KO^*;Cic*^KO^ (*top*), *Pten*^KO^*;Kmt2d*^KO^ (*center*), or *Pten*^KO^*;Zfhx3*^KO^ (*bottom*). IHC antibodies target the luminal epithelial marker cytokeratin 18 (CK18) and the basal epithelial markers cytokeratin 5 (CK5) and p63. Scale bars equal 100 mm. Additional tumors in **Supplementary Figure 9d**. (**e**) PAM50 prostate subtype scores for tumors with *TMPRSS2:ERG* fusion, SPOP mutation, ZFHX3 LOF, CIC mutation or deletion, KMT2D mutation, and TP53 mutation. (**f**) Comparison of putative driver mutation prevalence between castration-resistant SETs versus androgen-intact *Trp53*^KO^*;Msh2*^KO^ SETs. For each putative driver, the prevalence in castration-resistant SETs (*n* = 12) containing a point mutation is graphed versus the corresponding prevalence in androgen-intact SETs (*n* = 36), with jitter for visualization. (**g-k**) Kaplan-Meier inferred overall survival analysis from Caris Life Sciences cohort for CHEK1, KIFC1, RFC3, MDK, and ASPM stratified by gene expression quartile. Performance characteristics in **Supplementary Table 2**.

To investigate the biological basis of these survival associations, we characterized the lineage and hormone-response phenotypes associated with SET-identified alterations. Therapy-naïve Pten^KO^;Zfhx3^KO^ tumors exhibited high expression of luminal epithelial markers (CK18) and robust nuclear androgen receptor staining **(Figure 4d** and **Supplementary Fig. 9d–f**). By contrast, Pten^KO^;Kmt2d^KO^ and Pten^KO^;Cic^KO^ tumors exhibited high basal marker (Ck5, nuclear p63) expression and reduced nuclear AR levels^45,81–86^ (**Figure 4d, Supplementary Fig. 9d-f**). These findings indicate that SET-identified alterations correspond to lineage-associated hormone-response states that have historically been difficult to interrogate at genome scale. To extend these observations clinically, we analyzed transcriptomes from Caris-profiled patient tumors using PAM50 intrinsic subtyping^87,88^ (**Figure 4e**). *ZFHX3*-altered tumors exhibited a subtype distribution similar to that of *SPOP*-altered tumors, dominated by the Luminal A subtype and consistent with favorable clinical outcomes (**Figure 4e**). By contrast, *CIC*-altered tumors showed the highest Basal subtype prevalence, whereas *KMT2D*-altered tumors showed the highest proliferative Luminal B prevalence, consistent with their associations with poorer clinical outcomes (**Figure 4e**). Collectively, these findings indicate that alterations in *ZFHX3*, *CIC*, and *KMT2D* correspond to distinct evolutionary trajectories linking lineage identity, hormone response, and clinical outcomes in MMRP prostate tumors.

SETs identified therapy-sensitive clonal lineages even with minimal macroscopic tumor response, such as with TP53-altered tumors^5,61–64^ (**Fig. 3c-d, Supplementary Fig. 6b**), so we asked whether such SETs could also identify clinically relevant sensitization candidates, even among genes rarely mutated in patients. In androgen-intact Trp53^KO^;Msh2^KO^ SETs, genome-wide analysis again nominated pathogenic variants in *Zfhx3* alongside established hormone-response genes such as *Fgfr1* and the AR coactivator *Ep300*, both of which are being therapeutically targeted in combination with AR pathway inhibition^89–94^ (**Figure 4f**). Notably, androgen-intact SET evolution recurrently selected pathogenic variants in *ASPM*, *MDK*, *RFC3*, *CHEK1*, and *KIFC1*, genes spanning mitotic spindle regulation, DNA replication, and growth factor signaling^100–104^, despite their infrequent mutation in patient tumors. Transcriptional stratification of these gene candidates among Caris-profiled prostate tumors demonstrated that even modest reductions in expression from the highest expression quartile were associated with significant increases in inferred overall survival, often exceeding one year (**Figure 4h-l**). This is consistent with the selective enrichment of pathogenic variants in androgen-intact SETs, supporting a role for reduced gene activity in hormone therapy sensitization. Thus, SETs nominate these as candidate therapeutic targets for combination with AR pathway inhibition, analogous to efforts targeting *FGFR1* and *EP300*. These findings suggest that sparse mutational evolution can reveal clinically relevant therapeutic vulnerabilities even among genes that are infrequently mutated in patient tumors. Collectively, these findings demonstrate that by decoupling tumor evolution from chromosomal instability, SETs recover clinically relevant and actionable genetic and transcriptional determinants of therapeutic response that converge on distinct lineage-associated evolutionary states in cancer.

## Discussion

Interrogating the determinants of therapeutic response in vivo has been fundamentally limited by genomic complexity, in which chromosomal instability obscures gene-level inference. Here, we demonstrate that by separating tumor evolution from chromosomal instability, tumor evolution itself can be harnessed as a genome-scale evolutionary interrogation system. In this setting, stochastic mutation and physiological selection encode therapeutic fitness into sparse, interpretable genetic variation, enabling direct identification of the determinants of response. By initiating tumorigenesis from non-malignant substrates and constraining evolution to point mutations, this approach preserves native tissue context, lineage identity, and microenvironmental selection while enabling high-resolution inference of therapeutic evolution. Applied to prostate cancer, this strategy reveals in vivo determinants of therapeutic response that are otherwise inaccessible to conventional experimental approaches.

Existing genome-wide perturbation systems rely on predefined knockout libraries and are typically applied to transformed cells, limiting their ability to capture physiological selective pressures, allelic diversity, and non-coding regulatory evolution in vivo. Analyses of patient tumors interrogate naturally occurring evolutionary trajectories under therapy, but chromosomal instability obscures inference of specific variants through multigenic co-alteration. As a result, both perturbation-based and observational approaches incompletely resolve the lineage-associated evolutionary states that govern therapeutic sensitivity. In contrast, by decoupling tumor evolution from chromosomal instability, the evolution-driven strategy described here converts physiological selection into sparse, interpretable mutational landscapes that permit direct inference of variants associated with therapeutic response.

An important implication of this work is that the genetic determinants identified define stable lineage-associated phenotypic programs. In androgen-dependent prostate cancer, alterations such as *ZFHX3* loss specify a differentiated luminal hormone-responsive state associated with therapeutic sensitivity, whereas alterations enriched during castration-resistant evolution correspond to proliferative and basal features and reduced androgen receptor activity. These findings indicate that therapeutic response is governed by the selection of distinct evolutionary trajectories that couple lineage identity, cellular state, and hormone dependence. Such relationships have historically been difficult to interrogate genome-wide under physiological selection. By enabling in vivo selection across genetically diverse populations, this approach reveals functional links between genotype, lineage state, and therapeutic response that are otherwise inaccessible.

The interpretability of this approach derives from its emphasis on point-mutation–driven evolutionary trajectories, whereas evolutionary processes dominated by large structural variants may require complementary strategies. In this context, mismatch repair deficiency serves as an experimental lens to constrain genomic complexity, not a model of disease prevalence. The resulting sparse mutational landscapes are well-suited for computational inference, including deep learning–based prediction of variant function, and enable systematic integration with transcriptomic and phenotypic analyses. Because tumors evolve from common genetically defined origins under controlled in vivo conditions, this system permits direct comparison of evolutionary states across therapeutic contexts, which is difficult to achieve in patient cohorts. Here, SETs were developed in SCID hosts in order to easily isolate tumor-intrinsic evolutionary trajectories, but future adaptation to syngeneic immune-competent settings will enable direct interrogation of immune surveillance and adaptive immune selection during tumor evolution. More broadly, reconstructing tumor evolution without chromosomal instability establishes a generalizable strategy for genome-scale interrogation of therapeutic evolution across cancer types.

## Methods

### Organoid lines

Murine prostate organoids were isolated from the prostate of the Rosa26-Cas9 knockin (Strain 024858, JAX)^95^ or C57BL/6J (Strain 000664, JAX) mice, aged 10 weeks old. Rosa26-Cas9 knockin organoids (**Figures 2a-g, 3a-c,f-g, 4f, Supplementary Figs. 3c-e, 4, 6**) were infected with adenovirus-Cre to induce Cas9 expression, also resulting in EGFP expression (through the cas9-p2A-egfp linkage). C57BL/6J murine prostate organoids were used for the barcode lineage tracing experiments and for the gene validation experiments (**Figures 2h, 3d,h-j, 4d, Supplementary Figs. 5, 8a-c, 9d-f**). Prostate tissues were dissociated and expanded as 2D organoid-derived monolayers^96–102^ in Complete Mouse Organoid Medium (CMOM) and grown as described previously^30,102^ with the following modifications to CMOM: the addition of 10uM of Y-27632 dihydrochloride is constant and the concentration of Noggin and R-spondin 1 is 10% and 5% conditioned media respectively. Cells were passaged when 90% - 95% confluent by using TrypLE Express Enzyme (Gibco, 12605010). Cells were propagated in a Binder CO2 incubator controlled at 37°C and 5% CO2 and were confirmed to be free of mycoplasma using a Lonza detection kit (LT07-318).

### In vivo organoid allografts

For organoid allografts, 2 million organoid cells were injected subcutaneously on bilateral flanks of 8-10 weeks old C.B-17 SCID (CB17SC-M, Taconic or Strain 001303, JAX) or C57BL/6 (Strain 000664, JAX) males in 50:50 matrigel:media. Tumor growth was monitored weekly by caliper measurements. Volume was calculated as 0.5 * (length) * (width)^2^. For castration, mice were surgically castrated by bilateral orchiectomy (**Figures 3b-c, 3f-g, 4f, Supplementary Figs. 6a-c**) or chemically castrated by subcutaneous injection with 30 mg/kg degarelix (in 5% D-mannitol aqueous solution) every 4 weeks (**Figure 3d,h-j**). For ARPI, mice were treated every weekday with oral gavage or intraperitoneal injection of 10 mg/kg apalutamide or enzalutamide (in 5% DMSO, 1% CMC, and 0.1% Tween80) (**Figures 3h-i** and **3j**, respectively). Our intention-to-treat tumor volume thresholds to begin treatment with ARPI were 200 mm^3^ (**Figure 3b, Supplementary Figs. 6a-b**) and 50 mm^3^ (**Figure 3h-j**). Kaplan-Meier survival analysis was performed on GraphPad Prism using tumor kinetic data. For **Figures 3b, Supplementary Fig. 6b**, tumors in mice where the contralateral tumor size triggered ARPI treatment were censored at the time of said trigger. All animal care and experimental procedures were per the protocol approved by the University of California San Francisco Institutional Animal Care & Use Committee (#AN205866-00A).

### Ex vivo cell line isolation

Freshly dissected tumors of 50-100 mg were minced to approximately 2-3 mm^3^ pieces and digested with 5 ml of 2.4 mg/ml Collagenase A (Sigma, 10103576001) and 1 ml of 10 mg/ml DNase I (Sigma, 10104159001) in DMEM supplemented with 1% Pen/Strep and 0.5% Amphotericin B (Gibco, 15290018) at 37°C with agitation for 1 to 2 hours. Subsequently, 2 ml of TrypLE Express Enzyme (Gibco, 12605010) was added, and the digestion mixture was incubated at 37°C for 5 minutes. The digestion reaction was stopped with 10% FBS-containing DMEM and the cells were filtered with a 70 mm strainer. Cells were spun down at 200xg for 5 min, resuspended in CMOM and cultured as the organoids above.

### CRISPR/Cas9 Gene Editing

CRISPR/Cas9 gene knockout was performed either by introducing single-guide RNAs (sgRNA) cloned in pCRISPRia-v2 lentiviral vector (Addgene, plasmid #84832) in the Rosa26-Cas9 knockin organoids (**Figures 2a-g, 3a-c,f-g, 4f, Supplementary Figs. 3c-e, 4, 6**) or by electroporation of Cas9/sgRNA ribonucleoprotein (RNP) complexes in WT BL/6 organoids for barcode lineage tracing and gene validation experiments (**Figures 2h, 3d,h-j, 4d, Supplementary Figs. 5, 8a-c, 9d-f**). Lentivirally infected organoids were infected with the indicated lentivirus, selected with puromycin and further selected for BFP and GFP by FACS. For electroporation, we ordered either single-guide RNAs or Gene Knockout Kits, containing multi-guide RNA oligos (*n =* 3), from Synthego (now EditCo, California, USA) for each gene of interest (**Supplementary Table 3**). RNP complexes were prepared by combining 150 pmol of the multi-guide RNA oligos with 125 pmol of Alt-R Cas9 (IDT, 1081059) in PBS and incubated this at room temperature for 10 – 20 minutes. Subsequently, RNP complexes were added to 200,000 cells, resuspended in P1 buffer (Lonza, V4XP-1024), and cells were nucleofected using the 4D-Nucleofector Unit (Lonza Group AG, Switzerland) according to the manufacturer’s protocol. 10 minutes after electroporation, cells were gently resuspended in growth medium and seeded in 6-well plates. Successful gene knockout was confirmed by Western Blot (Methods: Immunoblotting) and Sanger Sequencing. For the latter, genomic DNA was extracted by resuspending the cells in 150 µL of QuickExtract (Lucigen, QE09050) and incubating in a thermal cycler at 65°C for 15 minutes followed by 95°C for 15 minutes. Target regions were amplified by a standard Polymerase Chain Reaction (PCR) with primers indicated in **Supplementary Table 3**. PCR products were column purified using the QIAquick PCR Purification Kit (Qiagen, #28106) and submitted for sequencing via the Sanger method (Azenta). The Synthego Inference of CRISPR Edits (ICE) tool was used to quantify knockout scores from the Sanger sequencing trace files ^103^.

### Barcode lineage tracing

CRISPR/Cas9 knockout of Msh6 was performed in C57BL/6 mouse-derived prostate organoid cells by electroporation of Cas9/sgRNA ribonucleoprotein complex alongside a non-targeting control (Methods: CRISPR/Cas9 Gene Editing). These lines were cultured for 8 months to allow for the accumulation of mutations in the MMRD line. The cells were then barcoded using an approach adapted from the ClonMapper system^48^, in which the barcoding plasmid backbone was substituted with CROP-Seq v2 vector (Addgene Plasmid #127458)^104^. As per the protocol, cells were infected with barcode-containing lentivirus at a low multiplicity of infection (MOI = 0.1) to ensure most cells incorporated only a single barcode. Barcoded cells were selected for with 5 µg/mL puromycin. After barcoding, cells were cultured for 26 passages to allow for the stabilization of the barcode system. Then, additional tumor suppressors (Pten or Trp53) were knocked out via RNP electroporation. Treatment arm mice began treatment on degarelix 2 weeks prior to organoid cell injection.

### In vitro growth assays

Murine prostate organoid cells were seeded in 96-well plates at a density of 250 or 125 cells/well. Cell growth was monitored by live-cell imaging every 4 to 6 hours, using an Incucyte S3 (Sartorius, USA). Phase masking was used to quantify the percentage of confluence in each well for each timepoint. For further analysis of the growth kinetics, a logistic growth curve was fitted (Python v3.12), using the following equation:

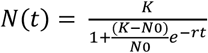

With N(t) the confluence at time t, N0 the initial confluence, K the carrying capacity and r the exponential growth rate. We restricted the model by setting K to 100, as this is the maximal confluence that can be reached in a well. To determine ARPI response in vitro, cells were treated with 10 µM enzalutamide (dissolved in DMSO) or 0.1% DMSO as experimental control.

### Immunoblotting

Cells or whole prostates, dissected from 10-12 week old C57BL/6 mice, were lysed in TNTE lysis buffer (50mM Tris, 150mM NaCl, 0.5% (v/v) Triton X-100, 1mM EDTA) containing 50mM NaF, 10mM Na4O7P2×10H2O, 10mM Na3VO4, 10mM PMSF and protease inhibitor cocktail (Sigma, P8340) as previously described ^105^. The prostate tissue was homogenized using a bead mill homogenizer (OMNI international) for 30s x 3 cycles. Total protein lysates were separated using either Nupage 7% Tris Acetate protein gels (Thermo Fisher), for Cic and Kmt2d immunoblotting, or Nupage 4-12% Bis-Tris protein gels (Thermo Fisher) for the rest of the proteins investigated. The proteins were transferred onto nitrocellulose membrane (Biorad) for 4h (for Cic and Kmt2d immunoblotting) or 1.5h for the rest. The blots were probed with primary antibodies as detailed in **Supplementary Table 4** and detected with SuperSignal Dura reagent (Thermo Scientific, 34075).

### Tumor Histology

Freshly dissected tissue was fixed in 10% formalin for 24 hours. After fixation, samples were stored in 70% ethanol at 4°C. Histology was carried out at Histowiz Inc. using a fully automated workflow and Standard Operating Procedures. Briefly, paraffin blocks were sectioned at 4μm thickness. The immunohistochemistry assay was performed on a Leica Bond RX automated stainer (Leica Microsystems). The tissue slides were dewaxed with xylene and alcohol-based solutions. Epitope retrieval was performed by heat-induced epitope retrieval (HIER) using a citrate-based pH6 solution for 20 minutes. After the primary antibody incubation (antibody details in **Supplementary Table 4**), samples with rabbit antibodies were processed with the Bond Polymer Refine Detection Kit, while mouse primary antibodies were incubated with Powervision Ms HRP as the secondary antibody (Bond Polymer Refine Detection Kit, Leica Microsystems). Then, samples were counterstained with hematoxylin according to the manufacturer’s protocol. The slides were dried, covered using the advanced TissueTek-Prisma Coverslipper, and scanned at 40X using a high-quality Leica Aperio AT2 slide scanner. Interpretation was provided by genitourinary pathologists. The AR score was determined by multiplying nuclear AR intensity (0-3+) by the percentage of AR+ cells (0-100%) to get a final score between 0 and 300.

### Bioinformatic analyses

#### Sample collection and whole-exome sequencing

Murine samples were collected from tumors, pregrafts and SELs to investigate genomic evolution. Genomic DNA was isolated from cell pellets or tumor tissue using the Qiagen DNeasy Blood & Tissue Kit (Cat. 69504). Whole-exome sequencing (WES) was performed by MedGenome, utilizing the Agilent SureSelectXT Mouse All Exon kit for library preparation. Sequencing was carried out on an Illumina NovaSeq platform, producing 100 bp paired-end reads at the mean depth of 52X. MedGenome processed the raw sequencing data, including quality control, read alignment, and base recalibration, before delivering the processed BAM files to our team for further analysis. The mouse reference genome assembly GRCm38 was used for all genomic and bioinformatic analyses.

#### Variant calling and initial filtration

Following receipt of the BAM files, we performed both germline and somatic variant calling on each sample using the Strelka2 (v2.9) variant caller^106^ configured with default parameters for whole exome to detect single-nucleotide variants (SNVs) and indels (1-49 base pairs). Only variants designated as “PASS” were retained. For somatic variant calling, pre-CRISPR normal wild-type murine prostate cells were used as the matched normal sample. To eliminate potential stromal or host-specific variants from our tumor samples (excluding pregraft samples), we conducted an additional filtration step. Variants from the tumor samples were compared to whole-exome sequencing data from the host mouse’s tail tissue, which was processed through the same pipeline as the tumors and pregrafts. For SNVs, we excluded all tumor variants that matched exactly with those found in the host tail tissue. For indels, however, we recognized that slight positional discrepancies could occur due to alignment uncertainties inherent in indel detection^107–109^. Therefore, we excluded indels if they overlapped with indels in the host mouse within a 10-bp window, using bedtools version 2.31.0^110^. By considering a 10 bp overlap, we ensured that host-specific indels with slight positional variations were excluded, preventing them from being misclassified as tumor-specific mutations. At this stage, we did not apply any filtration based on variant allele frequency or known SNPs in order to maximize the pool of potential variants for the downstream analyses. Following these filtering steps, the final variant set consisted of 1,261,865 mutations, including 869,189 SNVs and 392,676 indels.

#### CRISPR-targeted gene exclusion

In all CRISPR-engineered samples, including pregrafts, tumors, and ex-vivo cultures, variants within CRISPR-targeted genes were excluded altogether from further analysis to prevent any bias introduced by pre-existing genetic modifications. For instance, in tumors characterized by *Pten*^KO^*;Msh2*^KO^ genotype, all residual variants within Pten and Msh2 were systematically removed.

#### Gene mutation burden

We defined and calculated the gene mutation burden (GMB) for SNVs as the total number of coding SNVs (both nonsynonymous and synonymous) per megabase (Mb) across all tumors, adjusted for the gene length for each gene and further normalized by the number of samples. For the indel-specific GMB, we used the number of coding indels, including both frameshift and in-frame mutations, with the same normalization approach.

#### Copy Number Alteration and Fraction Genome Altered analyses

Copy number alterations (CNA) were assessed using CNVkit version 0.9.10^111^. CNVkit with default parameters calculated log2 copy ratios, analyzing tumor and pregrafts samples referencing normal murine prostate tissue to detect regions of chromosomal gains and losses.

Moreover, to quantify genomic instability in our samples, we calculated the Fraction of Genome Altered (FGA) based on log2 ratios. Briefly, a diploid baseline was assumed, where a single-copy gain corresponds to a log2 ratio of approximately 0.585 and a single-copy loss of -1.0. However, due to tumor heterogeneity and the presence of stromal cells, these values were adjusted to avoid underestimating CNAs^2^. A conservative threshold of ±0.2 in log2 scale was used to account for sample purity and clonality, ensuring detection of lower-amplitude CNAs. Specifically, this threshold corresponds to a copy ratio of 1.14 for gains and 0.62 for losses, approximately 24% below theoretical values, compensating for normal cell admixture. FGA was then calculated as the fraction of the genome showing gains or losses beyond these thresholds, providing a measure of genomic instability. However, due to the low frequency of copy number alterations observed across the tumors, as demonstrated by the FGA versus TMB analysis, subsequent analyses were predominantly focused on SNVs and indels.

#### Mutation categorization and enrichment

To thoroughly characterize the mutation landscape of each tumor and its corresponding pregraft, we performed pairwise comparisons of all mutations (SNVs and indels) between each tumor and its cognate pregraft. For each mutation, an exact match between the tumor and pregraft was required to confirm it as shared. To further categorize mutations, we assessed the variant allele frequency (VAF) of each mutation in both the tumor and its pregraft tissue. For mutations detected in the tumor but absent in the pregraft, we assigned an arbitrary VAF of 0.01 in the pregraft to facilitate consistent comparison across samples.

We classified tumor variants into three categories based on their comparison with mutations in the cognate pregraft sample:

I. SET-specific: mutations that were not detected in the cognate pregraft;
II. Shared, enriched in the SET: mutations detected in the cognate pregraft with a SET/pregraft VAF ratio of ≥ 2;
III. Shared, not enriched in the SET: mutations detected in the cognate pregraft with a SET/pregraft VAF ratio of < 2.

Similarly, we classified pregraft variants into three categories based on their comparison with mutations in the combined set of cognate tumors:

I. Shared, enriched in SET: mutations detected in at least one cognate tumor, with a SET/pregraft VAF ≥ 2;
II. Shared, not enriched in SET: mutations detected in at least one cognate tumor, with a SET/pregraft VAF < 2;
III. Pregraft-specific: mutations not detected in any cognate tumor.

### Functional prediction

Functional prediction of variants was performed using Ensembl Variant Effect Predictor (VEP) version 88^112^ and SnpEff version 5.0d^113^ independently. This analysis allowed us to identify variants predicted to have HIGH or MODERATE impact, which we prioritized for further investigation. The use of both VEP and SnpEff aimed to enhance confidence in the functional impact predictions by cross-validating results from two independent tools. We observed a high degree of concordance between VEP and SnpEff in predicting variant impacts. In instances of discrepancy between the tools, we followed the VEP prediction as the final assessment.

### Variant effect prediction upon histone modification/mRNA expression changes by AlphaMissense and AlphaGenome

To evaluate the functional consequences of both coding and noncoding DNA variants identified in the mouse samples, we employed a deep learning-based predictive framework utilizing AlphaMissense and AlphaGenome.^70,77^ Specifically, AlphaMissense was applied to assess the pathogenic potential and structural impact of missense variants within protein-coding regions. Concurrently, AlphaGenome was utilized to evaluate the regulatory implications of noncoding variants. By integrating these two predictive models, we systematically profiled how the identified genomic variants contribute to both protein dysfunction and gene regulatory perturbation in the mouse genome.

Because AlphaMissense provides pathogenicity predictions only for human proteins, mouse somatic missense variants were annotated by ortholog mapping prior to scoring. Variant calls in MAF format (mm10 coordinates) were lifted over to GRCm39 and mapped to their human orthologs using the h2m library (bioh2m), which performs codon-aware mouse-to-human variant modeling (m2h direction) against GENCODE references (mouse GRCm39/GENCODE M34; human GRCh38/GENCODE v44).^114^ For each mouse missense variant, h2m identified the orthologous human gene and canonical transcript, and generated the corresponding human protein substitution (human_HGVSp). Successfully modeled human variants were then queried against the precomputed AlphaMissense hg38 score table by exact genomic coordinate and allele using tabix, with the non-canonical isoform table queried as a fallback when no canonical-transcript hit was found. Mapped missense variants that could not be assigned a score on the first pass were re-run through the same mapping/lookup pipeline to recover additional scores, and any newly scored variants were patched back into the final annotated MAF. For the discovery analysis done here, a variant was treated as a potential pathogenicity event when its mapped AlphaMissense score was ≥ 0.5 (or it was a VEP HIGH-impact mutation, or an ESM-rescued AM-unclassified missense variant), restricted to variants with VAF ≥ 0.15.

#### ESM rescue of AlphaMissense-unclassified missense variants

Missense variants that the ortholog-mapping pipeline could not assign an AlphaMissense class (due to coordinate mismatch, failed mapping, lack of AlphaMissense score, or failed GRCm38 to GRCm39 liftover) were rescued using protein language-model scoring on the native mouse protein, bypassing the human-ortholog requirement. For each such variant, the mouse protein sequence was retrieved from GENCODE vM34 (gencode.vM34.pc_translations.fa) by transcript ID, the wild-type residue identity at the substituted position was verified, and a window of up to 1,022 amino acids centered on the variant was extracted. Each substitution was scored as a masked-marginal log-likelihood ratio (LLR = log P(alt) − log P(wt)) at the masked position, using two model families: ESM-2 (esm2_t33_650M_UR50D) and an ensemble of five ESM-1v models (esm1v_t33_650M_UR90S_1–5, averaged to give a mean ESM-1v LLR), run with fair-esm v2.0.0 (PyTorch 2.6, CUDA, NVIDIA L40S GPU).^115,116^ To place these LLRs on an interpretable scale, a calibration panel of AlphaMissense-classified missense variants from the same cohort (up to 1,000 each of likely-pathogenic, likely-benign, and ambiguous, sampled with fixed seed) was scored in the same run, and per-class LLR distributions were computed. A variant was flagged as “ESM-rescued” when it was an AlphaMissense-unclassified missense variant and its scores fell below both calibration-derived deleterious thresholds (mean ESM-1v LLR ≤ −6.38 and ESM-2 LLR ≤ −6.10, anchored to the likely-pathogenic calibration distribution); a stricter threshold set (≤ −6.66 and ≤ −7.5) was retained for sensitivity analysis.

### Noncoding variant prioritization with AlphaGenome

To nominate regulatory noncoding variants, cohort MAFs were filtered to VEP “Modifier” variants, and intersected with a regulatory BED was built as the union, over cancer panel genes described in following section, of TSS ± 2 kb windows (strand-aware, Ensembl r92 / GRCm38) and ENCODE SCREEN Registry V4 mm10 cCREs falling within ±50 kb of the gene body (66,012 merged intervals); coordinate-based attribution was used throughout to avoid gene-call misattribution. Variants overlapping any regulatory interval were retained as candidates (PTEN/MSH2, 1,759; P53/MSH2, 1,119 unique variants) and scored with the Google DeepMind AlphaGenome API (Python SDK v0.6.1, Organism.MUS_MUSCULUS, 18 recommended mouse scorers, 100 kb window centered on each variant). For each variant, predicted RNA-seq log₂ fold-changes (GeneMaskLFCScorer) on cancer-set genes within the window were summarized as the maximum |LFC| across biosamples, the maximum |per-gene median| across all 102 mouse RNA-seq tracks, and—the primary score—the maximum |mean LFC| over a curated 24-track epithelium-relevant subset (kidney, bladder, mammary, pancreas, ovary, testis, intestine/colon, lung, stomach, liver, adrenal), excluding hematopoietic, neural, muscle, adipose, and embryonic-stem-cell tracks. Because AlphaGenome’s mouse track set lacks purified prostate-epithelial RNA-seq, the epithelium-relevant subset served as the closest surrogate, and predicted effect sizes were treated as hypotheses for validation in accompanying RNA-seq. SET-derived mouse tumor samples were paired-end sequenced using the Illumina TruSeq platform. Raw reads were quality-trimmed using Trim Galore (v0.6.10), which utilizes CutAdapt (v1.18). Transcript abundances were quantified in Transcripts Per Million (TPM) using Salmon (v1.10.1) against the Ensembl GRCm39 reference transcriptome (release 108). Histone track modifications were also incorporated into the analysis to refine variant lists.

### Analysis of recurrent mutations

Moreover, while all genes were considered in the mutation pool, we chose to limit our attention to an aggregated list of 697 cancer-related genes. This list included pan-cancer genes provided by the UCSF500 and MSK-IMPACT panels, as well as significantly mutated prostate cancer genes identified in a meta-analysis of 1,013 patient tumors^117^ (**Supplementary Table 6**). Focusing on these genes ensured that our analysis was aligned with well-characterized oncogenic mutations relevant to human cancer.

### Mutational signature analysis

To systematically identify known mutational processes operative within our pregrafts and tumors, rather than extracting novel signatures *de novo*, we focused on fitting published MMRD and clock-like signatures. To this end, trinucleotide-inferred mutational signatures were calculated the using non-negative least squares method in MutationalPatterns version 3.2.0^118,119^ under R version 4.1.1. The contributions of signatures of interest were examined by assessing the relative contribution of each signature across samples. More specifically, we fitted known MMRD signatures (SBS6, SBS14, SBS15, SBS21, SBS26, and SBS44) and clock-like signatures (SBS1 and SBS5)^18,19^ to SNVs (and indels) to determine the degree of presence of these signatures in our engineered samples over time and across different genotypes. The analysis was performed on both the combined dataset and individual sample groups (pregrafts and tumors) to identify any group-specific signature patterns. We also enforced a maximum allowed reconstruction error of 0.2% between the observed and reconstructed mutation profiles, enhancing specificity by including only the most relevant signatures. Moreover, to avoid overfitting and the misattribution of signatures, we used the function fit_to_signatures_strict which employs an iterative approach where an initial fitting of all candidate signatures is performed, and then the signature with the lowest contribution is sequentially removed. At each iteration, the fit is recalculated, and the cosine similarity between the observed and reconstructed mutational profiles is assessed until the change between iterations exceeded a predefined cutoff. This method enhances specificity by preventing insignificant signatures from influencing the analysis.

### Quantification of Intratumor heterogeneity

Intratumor heterogeneity for tumors and pregrafts was quantified using the Shannon Diversity Index^120,121^ (SDI), which was calculated based on the distribution of VAFs within each sample:

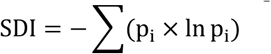

where 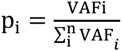.

### Clonality analysis and calculation of evolutionary distance

We inferred clonal architecture and phylogenetic relationships among our samples using CloneFinder, a computational tool designed for clonal inference from deep sequencing data^69^. CloneFinder reconstructs the phylogenetic tree of clones by leveraging parsimony-informative SNVs, enabling the identification of subclonal cell populations and their evolutionary relationships. To enhance the accuracy of clonal frequency estimates and minimize potential noise, we applied stringent filtering criteria to the SNVs. Specifically, we excluded SNVs with a total read depth less than 40 reads and those with fewer than 6 supporting alternative read counts, as has been successfully implemented in previous studies^122,123^. This filtering ensured that only high-confidence SNVs were included in the analysis, reducing the impact of sequencing errors and low-frequency artifacts. In this analysis, “clonal frequency” refers to the estimated proportion of cells within a sample that belong to a particular clonal population or subclone. We set a cutoff of 7.5% for estimated clonal frequencies to delineate significant clones, effectively excluding subclonal populations below this threshold due to potential limitations in detection sensitivity. An important assumption of CloneFinder is that the input data are free from CNAs, as CNAs can distort allele frequencies and hinder accurate clonal inference based solely on SNVs. As mentioned above, our samples (both tumors and pregrafts) exhibited minimal CNA events, allowing us to satisfy this assumption and proceed without incorporating CNAs into the clonality inference. This approach facilitated a clear resolution of clonal populations based exclusively on SNV data. The resulting phylogenetic trees were further analyzed using the ape package (v5.6.2) and ggtree (v3.0.4) in R v4.1.1 to extract evolutionary distances between clones. For tumors that appeared monoclonal, each clone represented a distinct tumor sample. Evolutionary distances were calculated using cophenetic distances derived from the phylogenetic trees, providing quantitative measures of genetic divergence among tumor subpopulations. These distances offered insights into the evolutionary dynamics and heterogeneity within and between tumors.

### Sample collection and barcode sequencing

Genomic DNA from the resulting tumors was isolated and prepared for sequencing using a 2-step PCR protocol to introduce sequencing adapters and unique dual indices, adapted from the ClonMapper system^48^ (**Supplementary Table 5**). Amplicon sequencing was performed on a NovaSeq X (UCSF CAT) with a target read depth of ∼10 million per sample. Limit of detection was imposed by input gDNA amount – 1µg used which corresponds to ∼150,000 diploid genomes or ∼0.001%. Analysis of amplicon sequencing was performed using PyCashier v23.1.2 with a count filter of 20 and default parameters^48^ (<u>GitHub</u>). The frequency of each barcode was then calculated as the fraction of total PyCashier-filtered reads corresponding to each unique barcode.

### Quantification of heterogeneity

For ClonMapper barcoded tumors, Shannon diversity was calculated analogously to above, using the frequency of each barcode (BC) within a tumor, i.e.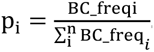.

Barcoding also affords the ability to examine inter-tumoral heterogeneity by comparing the dissimilarity of tumor barcode composition. For this, we calculated the Bray-Curtis dissimilarity for pairs of samples:

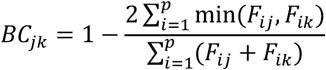

where *F_ij_* is the frequency of barcode i in sample j, *F_ik_*is the frequency of barcode i in sample k, and p is the total number of barcodes in the samples.

### Analysis of patient data

#### Patient datasets

See **Data availability** for dataset sources. Genomic alterations, including mutations, copy-number alterations and structural variations, and tumor mutation burden, were gathered from the cBioPortal-preprocessed data. Microsatellite instability of the MSK-MetTropism dataset was calculated by MSISensor^124^, and we rephrased MSI Stable, Indeterminate, and Instable as MSS, intermediate, and MSI-H, respectively. Regarding the Caris cohort, next-generation sequencing (592-gene or whole exome) was performed for prostate adenocarcinoma patient samples submitted to Caris Life Sciences (Phoenix, AZ). Patients were stratified by the presence of pathogenic/likely pathogenic mutations (interpreted by board-certified molecular geneticists and categorized according to the American College of Medical Genetics and Genomics standards), homozygous deletions, gene expression and for ZFHX3, the presence of any truncating mutation upstream of codon 3381^125^. We classified all patients analyzed as Microsatellite stability by NGS^126,127^. Real-world overall survival was obtained from insurance claims data and calculated as the time from biopsy to time of last follow-up or death. Hazard ratio (HR) was calculated using the Cox proportional hazards model, and P values were calculated using the log-rank test. Treatment includes any of leuprolide, degarelix, relugolix, enzalutamide, apalutamide, darolutamide, and/or abiraterone.

#### PAM50 Analysis

The PAM50 signature classifies tumors into five subtypes - Normal, Basal, Her2-enriched, Luminal A and Luminal B-for samples with evaluable RNA expression for a list of 50 genes (ACTR3B, ANLN, BAG1, BCL2, BIRC5, BLVRA, CCNB1, CCNE1, CDC20, CDC6, NUF2, CDH3, CENPF, CEP55, CXXC5, EGFR, ERBB2, ESR1, EXO1, FGFR4, FOXA1, FOXC1, GPR160, GRB7, KIF2C, NDC80, KRT14, KRT17, KRT5, MAPT, MDM2, MELK, MIA, MKI67, MLPH, MMP11, MYBL2, MYC, NAT1, ORC6, PGR, PHGDH, PTTG1, RRM2, SFRP1, SLC39A6, TMEM45B, TYMS, UBE2C, UBE2T). To adapt the PAM50 signature to Prostate Cancer, the normal subtype was excluded.

#### Landscape of genomic alterations from cancer patients

The frequency of nonsynonymous SNVs, INDEL, amplification, and homologous deletion was collected from the TCGA^128^ and PCF/SU2C East Coast Dream Team^5^ in the cBioPortal for Cancer Genomics. All frequencies from protein-coding genes were aligned in the order of their positions in the chromosomes.

#### High-TMB / MSI-High exclusive mutations in co-localized genomic alterations

Two MMRD-predominant cancers, colorectal and endometrial cancer, using datasets from TCGA PanCancer Atlas^128^ were analyzed for differences in co-localized genomic alterations between MSS and MSI-High cancer patients and other cancers, melanoma and bladder cancer, were analyzed between high- and low-tumor mutation burden (TMB) cancer patients. Patients were in high- and low-TMB groups based on a cutoff of 10 nonsynonymous SNVs/Mb (NS/Mb)^16^ and in MSS and MSI-High groups based on cutoffs of MSISensor score 4 and 10 to be calculated the numbers of mutations and copy-number alterations in both groups. We defined high-TMB or MSI-High exclusively mutated, co-localized genes under these criteria: ≥ 5 patients mutated in high-TMB or MSI-High, ≤ 5 patients mutated in low-TMB or MSS, and ≥ 5 patients copy-number altered in low-TMB or MSS in cytoband-level. For the average altered patient percentage, the number of patients with genomic alterations in the selected genes was averaged per cytoband and divided by the total number of patients in each corresponding patient group. We graphed the cytobands that satisfied three or more selected genes per cytoband and satisfied less than 5% of average patients in both amplification and deletion at high-TMB or MSI-High.

### Comparison of gene variants between MSI-High and MSS Patient Tumors

#### MSI Enrichment Ratio

We compared gene variants between MSI-High and MSS patients at the level of individual genes. Patient tumor data were gathered from MSK-MetTropism^33^ based on MSISensor score^124^. For each gene, the frequency of variants (i.e., mutations, copy-number alterations, and structural variants) was calculated among MSI-High and MSS patients. To adjust the effect of mutation number, we calculated a log-transformed, TMB-normalized gene alteration ratio between MSI-High and MSS as:

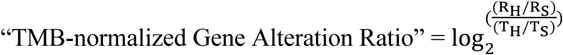

where RH is the altered patients’ ratio in MSI-High, RS is the altered patients’ ratio in MSS, TH is the averaged TMB in MSI-High, and TS is the average TMB in MSS. For the histogram, we filtered out genes that were altered in five or fewer patients.

#### GSEA

We also performed Gene Set Enrichment Analysis (GSEA^23,24^ to quantify gene set-level similarity between MSS and MSI-High by using the GSEApy package^129^ with GO.BP^130^, Reactome^131^, WIkiPathways^132^, and KEGG^133^. The inputs are the frequency of gene variants among MSS and MSI-High patient tumors. The input gene lists of MSS and MSI-High were applied to the prerank module for GSEA using the following parameters: maximum 500, minimum 5, and permutation number 1000.

### Statistical Analyses

For statistical analyses, R (version 4.1.1) and GraphPad Prism Software (version 10) or Python (version 3.12) were used. Different statistical tests were used depending on the type of data that was handled, including log-rank tests for survival analyses, mixed effects models for observing treatment effect on tumor volume over time, t-tests or ANOVAs (with post-hoc comparisons) for comparing the means of two or more groups, and binomial or hypergeometric tests for comparing population frequencies. Multiple hypothesis testing was accounted for where appropriate. These details are further specified in every figure caption. Tests were two-sided unless otherwise noted. p-values smaller than 0.05 were considered statistically significant. Mouse graft experiments consisted of a minimum of five tumors per condition. The sample size estimate was based on our experience with previous experiments^9^. No formal randomization process was used to assign mice to a given organoid injection, and experimenters were not blinded.

## Data and code availability

Publicly available patient datasets were collected from the cBioPortal for Cancer Genomics (https://www.cbioportal.org/)^134–136^: MSK-MetTropism^33^, NCI Patient-Derived Models Repository, and TCGA PanCancer Atlas studies^128^ were used for pan-cancer level analysis. For metastatic prostate adenocarcinoma, we used the dataset from PCF/SU2C East Coast Dream Team^5^. All high-throughput Whole Exome Sequencing (WES) data and raw barcode sequencing data from murine prostate tumors and organoids are available through NCBI’s Sequence Read Archive (SRA PRJNA1226760). WES analysis pipeline implementation available upon request.

## Acknowledgments

We are grateful to the cancer patients whose profiling and contributions made this work possible. We thank the Bose lab for valuable critiques. We further appreciate the discussions with many at UCSF, primarily through our departments, the UCSF Benioff Initiative for Prostate Cancer Research (Dr. Franklin Huang and Dr. Ross Okimoto), and the UCSF Bakar Aging Research Institute. We thank the editing contributions made to this manuscript by The Science Editors and the Arc Institute. The sequencing was performed at the UCSF CAT, supported by UCSF PBBR, RRP IMIA, and NIH 1S10OD028511-01 grants. This work was assisted by the UCSF Center for Advanced Technologies and is supported by the Diabetes Center at UCSF. Schematics were created in BioRender. R.B. is supported by National Institutes of Health grants DP2-CA290244 and K08CA226348, American Cancer Society grant RSG-22-021-01-CDP, US Department of Defense grants PC220482 and PC141542, Prostate Cancer Foundation grant 16YOUN01, the Benioff Initiative for Prostate Cancer Research, and the Bakar Aging Research Institute. M.J.R. was supported by National Institutes of Health grant F31CA278406. W.S. is supported by Prostate Cancer Foundation grant 22YOUN10. F.Y.F was supported by the Benioff Initiative for Prostate Cancer Research. H.G. is supported by the Arc Institute.

## Author contributions

R.B. conceptualized the study. R.M.B, M.J.R., W.S., S.B.H., V.L., A.A.C, L.H., and P.J. developed the methodology. M.J.R., S.B.H., V.L., A.A.C., L.H., J.C., Y.S.L., K.H., C.C.D., B.A.S., P.J., T.N., X.F., M.H., and A.N. conducted the experiments featured here. R.M.B., W.S., S.B.H., L.H., B.D., D.B, and S.W. performed analysis. M.J.R., W.S., S.B.H., V.L., L.H., K.H., and B.D. visualized the data. C.L.S., F.Y.F., H.G., and R.B. acquired funding for the project. R.B. administered the project. W.R.K., C.L.S., H.G., and R.B. provided resources. C.L.S. and R.B. provided supervision. R.M.B., M.J.R., W.S., S.B.H., V.L., L.H., and R.B. wrote the manuscript. We have a U.S. patent application related to this work (63/765,931).

## Materials & Correspondence

Address requests to Rohit Bose.

