## Supplementary Figures for "Reconstructing tumor evolution without chromosomal instability reveals in vivo determinants of therapeutic response"

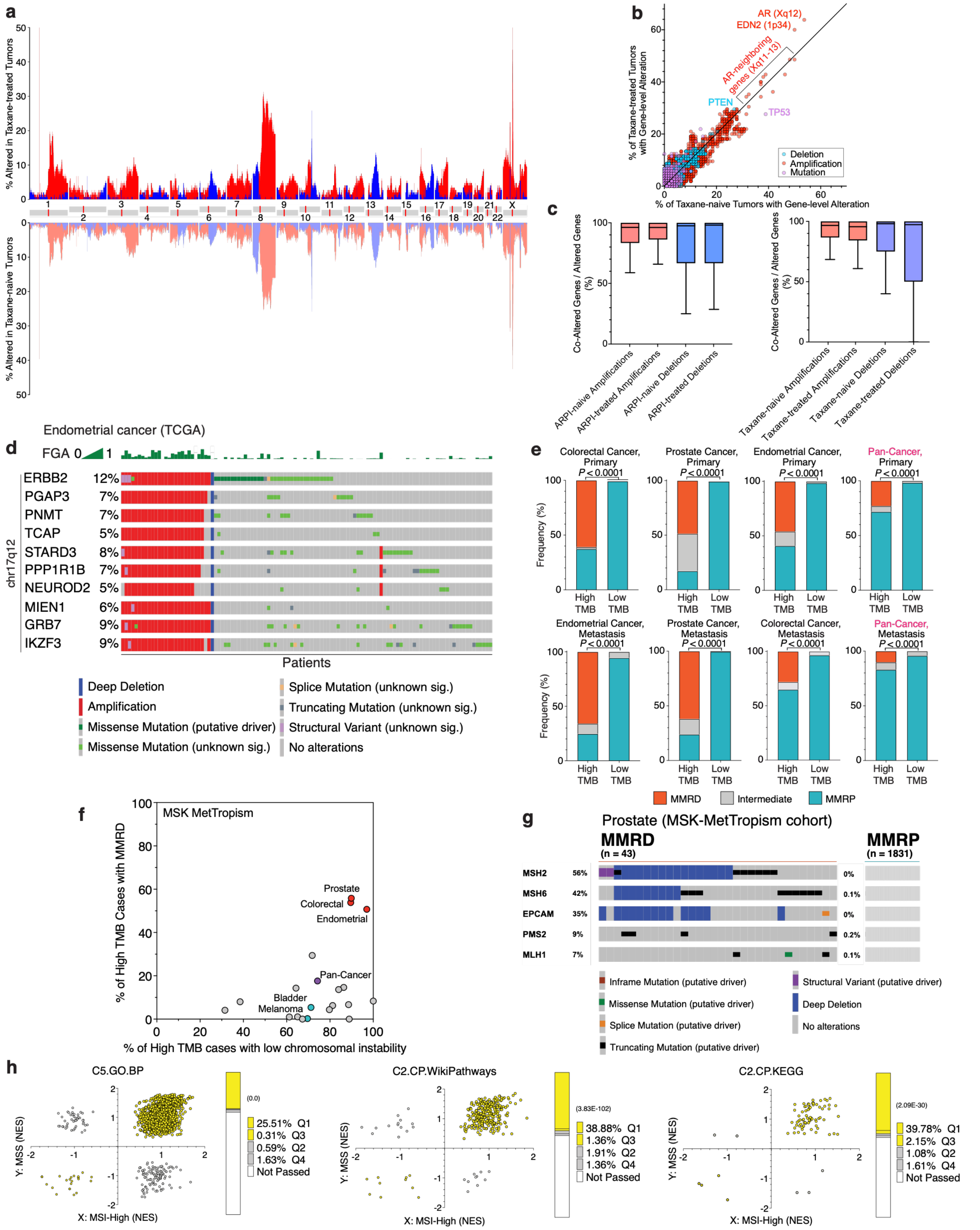


**Supplementary Figure 1 – Mismatch repair deficiency enables high-resolution inference of tumor evolution obscured by chromosomal instability**

(**a**) Landscape of copy number alterations in taxane-treated (*top*) and taxane-naïve (*bottom*) advanced prostate cancer patient tumors. The frequency of copy number alterations is graphed versus chromosomal locus. Copy number alterations are amplifications (red) and deep deletions (blue)[2]. Examples of OncoKB annotated cancer-related genes[137,138] are highlighted at loci subject to frequent copy number alterations. Whole exome sequencing (WES) results are from the PCF/SU2C East Coast Dream Team metastatic castration-resistant prostate cancer cohort[5] (*n* = 444). (**b**) The rate of deletion (blue), amplification (red), and mutation (magenta) of genes is compared in taxane-treated tumors versus taxane-naïve tumors in ARPI-treated patients within the PCF/SU2C East Coast Dream Team metastatic castration-resistant prostate cancer cohort[5] (*n* = 444). (**c**) For each chromosome, the number of copy number altered genes that are adjacent to another copy number altered gene is divided by the total number of copy number altered genes. This was then averaged across all chromosomes, and the value was compared between (*left*) ARPI-treated and naïve tumors and (*right*) taxane-treated and naïve tumors for amplifications (red) and deletions (blue). Tumors from the PCF/SU2C East Coast Dream Team metastatic castration-resistant prostate cancer cohort[5] (*n* = 444). (**d**) Point mutation pattern in cancer patients with high FGA tumors (mostly on the left) compared to low FGA tumors. The OncoPrint is derived from endometrial cancer patients (*n* = 529) in the TCGA cohort WES[128] obtained from cBioPortal[134–136]. (**e**) Relationship between microsatellite instability and high-TMB across primary (*top*) and metastatic (*bottom*) patient tumors. Frequency of MSI-High (High Microsatellite Instability) and MSS (Microsatellite Stable) among High-TMB (≥10 NS SNVs/Mb) and Low-TMB cases. Microsatellite Instability categorized using MSIsensor score and color-coded: red (≥10, MSI-High), grey (between 4-10, intermediate), or blue (≤4, MSS). A one-sided hypergeometric test was performed to compare the percentage of MSI-High tumors between groups. Data from MSK MetTropism cohort. (**f**) Cancer-specific relationship between frequency MSI-high cases versus low FGA cases among high-TMB (TMB ≥ 10 NS SNVs/Mb) patient tumors. Targeted panel sequencing from MSK-MetTropism cohort[33]: endometrial (*n* = 366), prostate (*n* = 79), colorectal (*n* = 595), small bowel (*n* = 18), non-melanoma skin (*n* = 79), hepatobiliary (*n* = 44), cervical (*n* = 25), anal (*n* = 16), head and neck (*n* = 74), pan-cancer (*n* = 24,753), pancreatic (*n* = 39), bladder (*n* = 459), melanoma (*n* = 543), esophagastric (*n* = 55), non-small cell lung (*n* = 1115), breast (*n* = 129), ovarian (*n* = 26), and small cell lung (*n* = 108). (**g**) Cancer-specific patterns of mutations that cause mismatch repair deficiency (MMRD). OncoPrints[135] of canonical MMRD genes[144–146] for MSI-High and MSS patient tumors. EPCAM is included as deep deletions of its polyadenylation site lead to transcriptional read-through, hypermethylation, and silencing of the neighboring MSH2 promoter[26]. Targeted panel sequencing from MSK-MetTropism cohort[33] prostate tumors: MSI-High cases (*n* = 43) and MSS cases (*n* = 1831). (**h**) Comparison of affected pathways in MSS versus MSI-High patient tumors. Each dot represents a pathway. For each pathway, the normalized enrichment score (NES) derived from gene set enrichment analysis (GSEA) of the gene variant frequency among MSS prostate cancer patient tumors was graphed versus that among MSI-High tumors. Positive or negative NES indicates that more or fewer patients, respectively, have gene variants in each corresponding gene set. Yellow fill highlights altered pathways located in the first and third quadrants. P-value by one-sided hypergeometric test. Targeted panel sequencing from MSK-MetTropism was analyzed with MSS (*n* = 1831) and MSI-High (*n* = 43) prostate cancer patients.

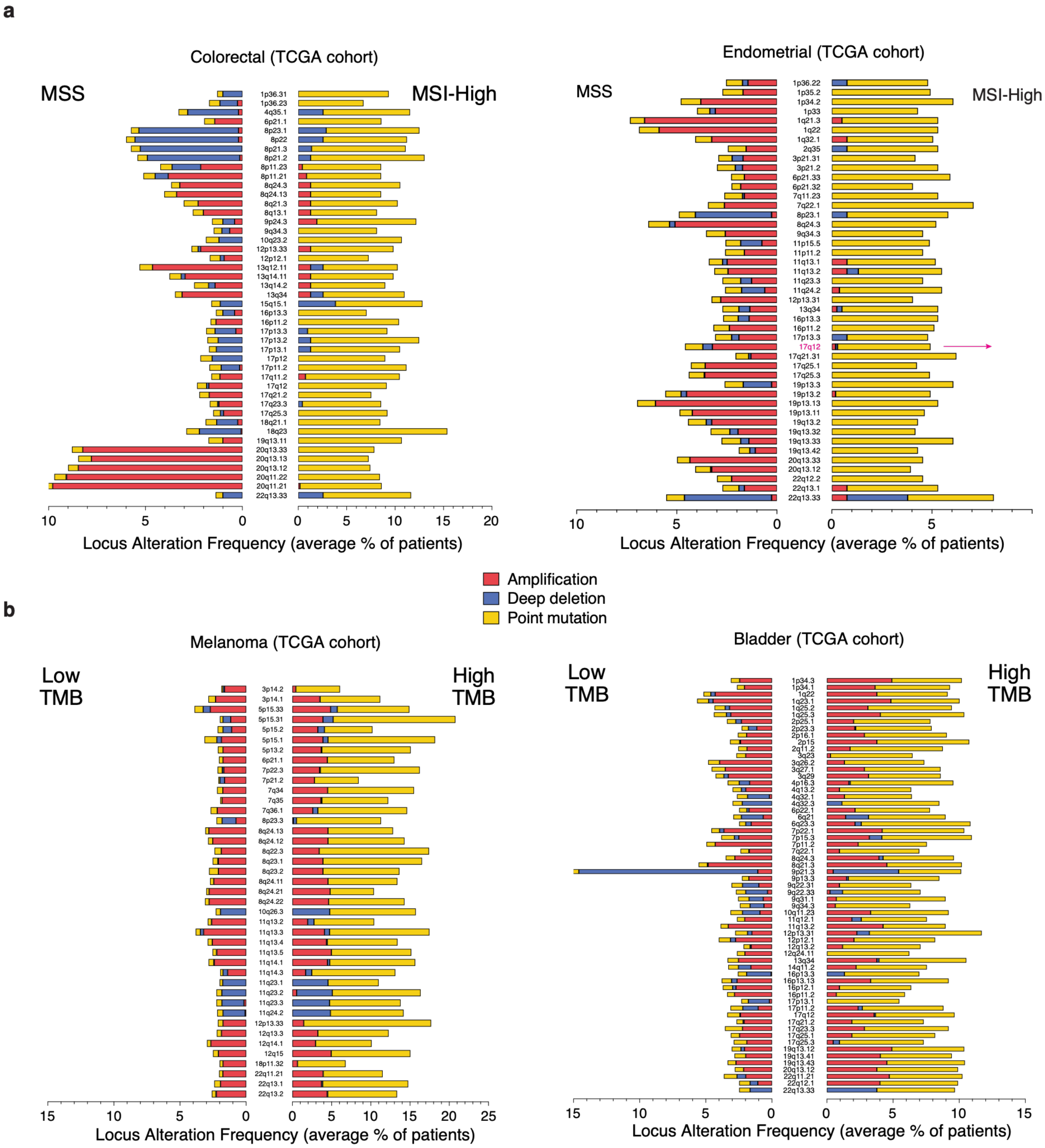


**Supplementary Figure 2 – Mismatch repair deficiency uniquely shifts recurrent tumor evolution from chromosomal instability to point mutational evolution**

(**a**) Comparison of gene variant patterns at chromosomal regions between MSS and MSI-High patient tumors. Cytobands are graphed versus the frequency of gene variant types indicated in red (Amplification), blue (Deep deletion), and yellow (Point mutation)[2]. WES from the TCGA cohort[128] was analyzed with (*left*) MSI-High ($n$ = 78) and MSS ($n$ = 496) colorectal cancer patients and (*right*) MSI-High ($n$ = 132) and MSS ($n$ = 368) endometrial cancer patients. (**b**) Comparison of gene variant patterns at chromosomal regions between Low-TMB (<10 NS SNVs/Mb) and High-TMB (≥10 NS SNVs/Mb) patient tumors. Cytobands are graphed versus the frequency of gene variant types indicated in red (Amplification), blue (Deep deletion), and yellow (Point mutation). WES from the TCGA cohort was analyzed for melanoma cases (*left*, Low-TMB ($n$ = 286) and High-TMB ($n$ = 154)) and bladder cases (*right*, Low-TMB ($n$ = 303) and High-TMB ($n$ = 106)).

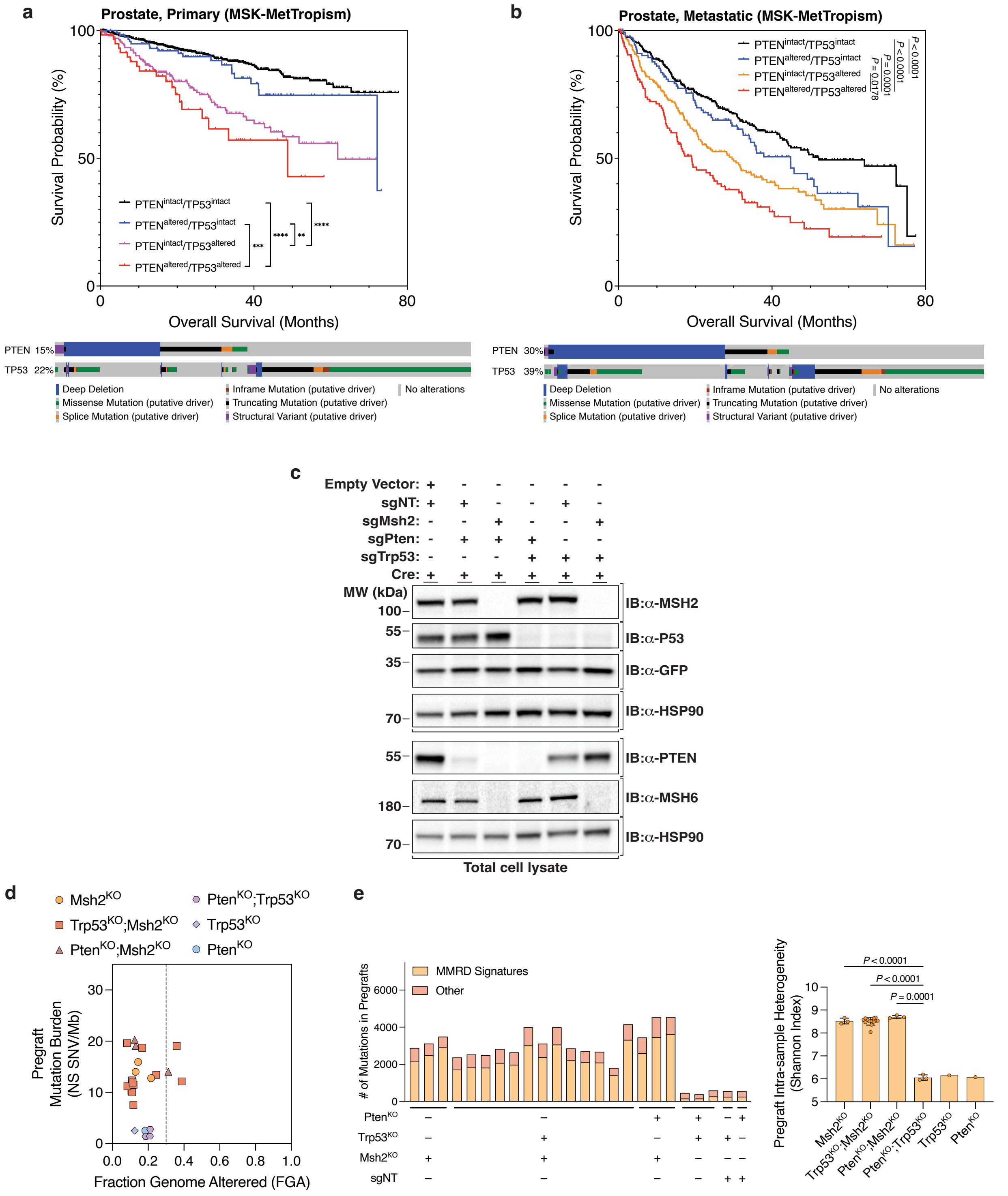


**Supplementary Figure 3 – MMRD pregrafts generate clinically relevant mutational heterogeneity without chromosomal instability**

(**a**) Clinical association of *PTEN*-alterations or *TP53*-alterations with patients' overall survival with primary site prostate profiled samples. *Top*: Kaplan-Meier survival probability curves for prostate cancer patients with primary site biopsies. *Bottom*: OncoPrint for putative driver alterations of *PTEN* and *TP53* in the primary prostate cancer patients corresponding to the top Kaplan-Meier curves. Data from MSK-MetTropism ($n$ = 1307). P-values by Log-rank test. (**b**) *Top*: Kaplan-Meier survival probability curves for prostate cancer patients with metastatic biopsies. *Bottom*: OncoPrint[134] for putative driver alterations in *PTEN* and *TP53* in metastatic prostate cancer patients. Data from MSK MetTropism[33] ($n$ = 859). P-values by Log-rank test. (**c**) Western blot shown of total cell lysates. Corresponding scheme in **Figure 2a.** (**d**) Analysis of mutation burden versus fraction genome altered for MMRD and mismatch repair proficient (MMRP) pregrafts. (**e**) *Left:* Mutational signature analysis of *Msh2*$^{KO}$-containing pregrafts. The total number of mutations attributed to an MMRD signature (SBS 6, 14, 15, 21, 26, or 44) was graphed as was that attributed to the clock-like (SBS 1 and 5) signatures[18,19]. *Right:* Shannon Diversity Index[120,121] calculated from point mutations. The same samples are included as **d**. P-values by one-way Welch's ANOVA with Dunnett's T3 multiple comparisons test.

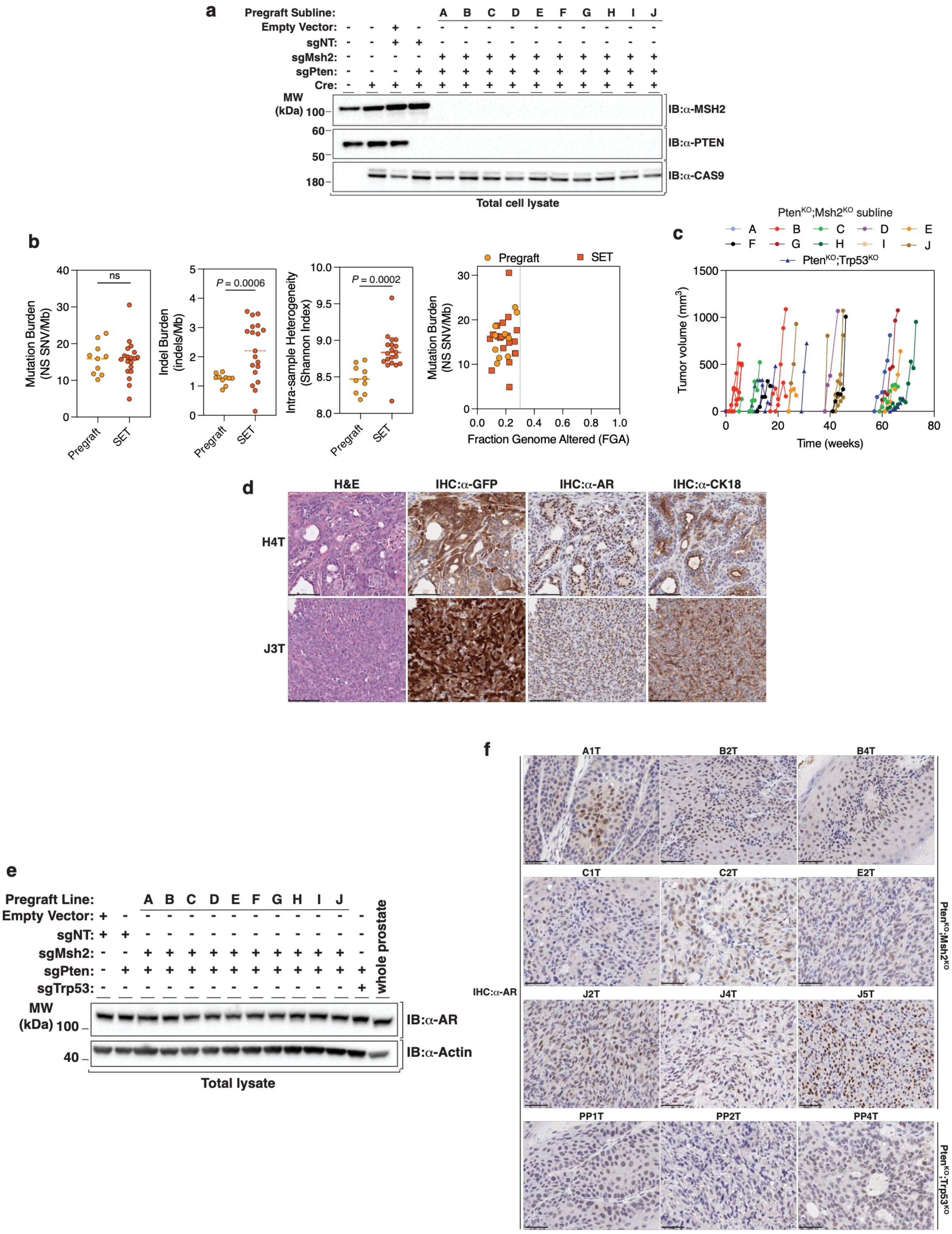


**Supplementary Figure 4 – SETs generate endogenous phenotypic heterogeneity during in vivo tumor evolution**
(**a**) Evaluation of MSH2 and PTEN protein levels via Western blot of total cell lysates from *Pten*$^{KO}$*;Msh2*$^{KO}$ (sgPten+sgMsh2) A–J pregraft sublines. (**b**) Mutation burden, indel burden, intra-sample heterogeneity (by Shannon Diversity Index), and fraction genome altered (FGA) for pregraft (*n* = 10) and SETs (*n* = 19). (**c**) Growth kinetics of

SETs arising from different MMRD clones. Tumor volume kinetics measured following murine injection of *Pten*$^{KO}$;*Msh2*$^{KO}$ A–J MMRD sublines (separated 44 days post-CRISPR) as well as MMRP *Pten*$^{KO}$;*Trp53*$^{KO}$ (sgPten+sgTrp53) or *Pten*$^{KO}$ (sgPten+sgNT) control lines ($n$ = 5 mice per subline or control lines). Injection of *Pten*$^{KO}$ pregrafts did not result in tumor formation by 72-week endpoint.(**d**) Two SETs (H4T, J3T) derived from 2 *Pten*$^{KO}$;*Msh2*$^{KO}$ sublines (H, J) have been subjected to hematoxylin and eosin (H&E) staining, as well as immunohistochemistry (IHC) for Green Fluorescent Protein (GFP) from the organoid pregraft line and two classic prostate tumor markers: Androgen Receptor (AR), and the luminal epithelial marker cytokeratin 18 (CK18). Scale bars equal 100 mm. (**e**) Nuclear hormone receptor protein levels among MMRD pregrafts compared to whole prostate. Western blot of Androgen Receptor (AR) and Actin in *Pten*$^{KO}$;*Msh2*$^{KO}$ A–J pregraft sublines versus MMRP controls and the whole prostate of a 12-week-old C57BL/6 mouse. (**f**) Representative immunohistochemistry (IHC) for the androgen receptor (AR) is shown for 9 distinct SETs (A1T, B2T, B4T, C1T, C2T, E2T, J2T, J4T, J5T) (in addition to **Figure 2e**) arising from *Pten*$^{KO}$;*Msh2*$^{KO}$ A, B, C, E, and J subline injections. IHC was also performed for 3 tumors (PP1T, PP2T, PP4T) arising from control *Pten*$^{KO}$;*Trp53*$^{KO}$ injections. Scale bars equal 50 mm. Pathologist quantitation in **Figure 2f**.

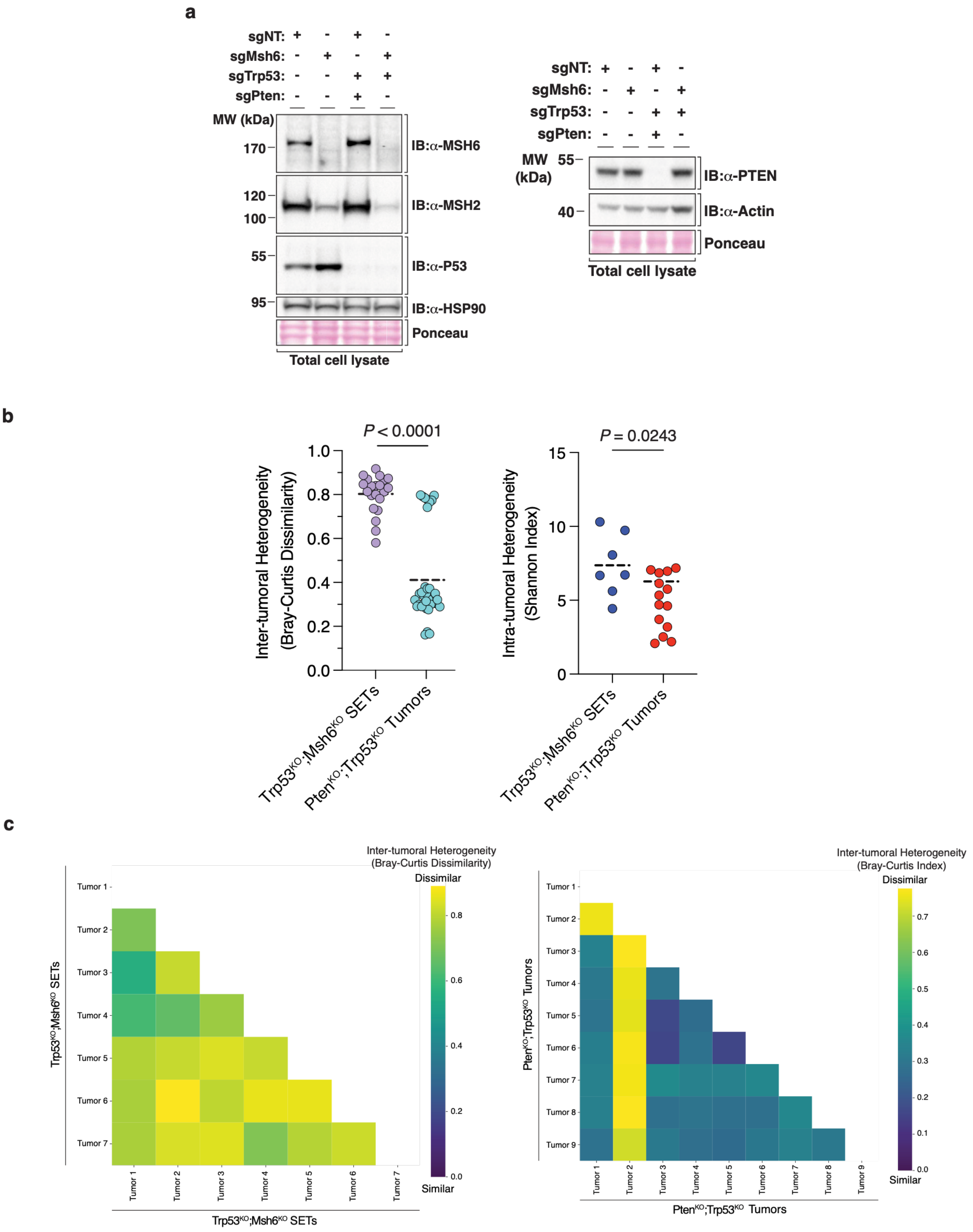


**Supplementary Figure 5 – SETs preserve stochastic clonal heterogeneity and avoid clonal dominance characteristic of conventional tumor models**

(**a**) Evaluation of MSH2/MSH6 heterodimer components following the introduction of sgMsh6, sgPten, sgTrp53 CRISPR guides. Total cell lysate of barcoded murine prostate organoid lines subjected to Western blot and Ponceau

S analysis. **(b)** *Left:* Comparison of inter-tumoral heterogeneity among SETs versus among MMRP tumors. Inter-sample mutational heterogeneity was assessed by Bray-Curtis Dissimilarity[139]. Plot shows all pairwise comparisons within the *Trp53*$^{KO}$;*Msh6*$^{KO}$ SET and *Pten*$^{KO}$;*Trp53*$^{KO}$ tumor groups, respectively. P-value by Welch's unpaired t-test. Barcode sequencing from *Trp53*$^{KO}$;*Msh6*$^{KO}$ SETs (*n* = 7) and *Pten*$^{KO}$;*Trp53*$^{KO}$ control tumors (*n* = 9). *Right:* Comparison of intra-tumoral heterogeneity among SETs versus among MMRP tumors. Intra-sample mutational heterogeneity was assessed by the Shannon Index[120,121] and calculated for each barcoded *Trp53*$^{KO}$;*Msh6*$^{KO}$ SET and MMRP *Pten*$^{KO}$;*Trp53*$^{KO}$ tumor. P-value by Welch's unpaired t-test. (**c**) Comparison of inter-tumoral heterogeneity among SETs versus MMRP tumors. Inter-sample mutational heterogeneity was assessed by the Bray-Curtis Dissimilarity[139]. Matrices show individual pairwise comparisons within the *Trp53*$^{KO}$;*Msh6*$^{KO}$ (sgTrp53+sgMsh6) SET group (*left*) and *Pten*$^{KO}$;*Trp53*$^{KO}$ (sgPten+sgTrp53) MMRP tumor group (*right*). Samples analyzed by barcode sequencing: *Trp53*$^{KO}$;*Msh6*$^{KO}$ SETs (*n* = 7) and *Pten*$^{KO}$;*Trp53*$^{KO}$ control tumors (*n* = 9).

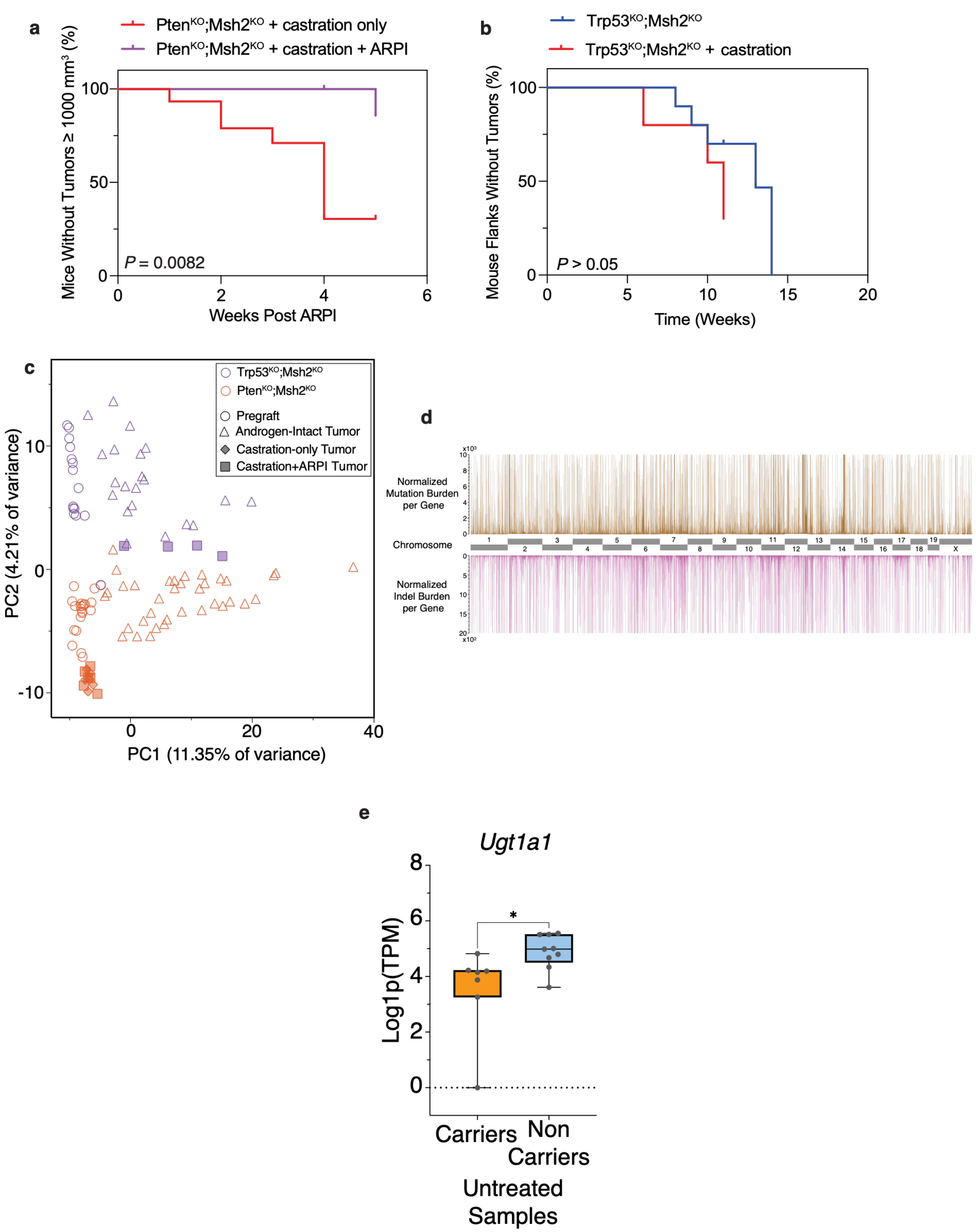


**Supplementary Figure 6 – SETs enable genome-wide interrogation of coding and non-coding determinants of therapeutic evolution**

(**a**) Effect of ARPI treatment on SET growth kinetics. Kaplan-Meier of mice without tumors ≥ 1000 mm$^3$ for *Pten*$^{KO}$;*Msh2*$^{KO}$ SETs upon ARPI treatment (apalutamide 10mg/kg). Castration-alone mice $n$ = 14, castration +

ARPI *n* = 6. P-value by log-rank test. (**b**) Kaplan-Meier of mouse flanks without tumors (no tumor ≥ 200 $mm^3$) of *Trp53*$^{KO}$*;Msh2*$^{KO}$ (sgTrp53+sgMsh2) SETs upon castration. Data from androgen-intact mice (*n* = 5) and castrated mice (*n* = 5). Comparison assessed by log-rank test. (**c**) Effect of cancer therapy and truncal genotype on SET mutational composition. Principal component analysis (PCA) of all SNVs and indels derived from all generated SETs and corresponding pregrafts. For *Pten*$^{KO}$;*Msh2*$^{KO}$ genotype: pregrafts (*n* = 21); androgen-intact SETs (*n* = 36); castration-alone SETs (*n* = 6); and castration + ARPI SETs (*n* = 6). For *Trp53*$^{KO}$;*Msh2*$^{KO}$ genotype: pregrafts (*n* = 15); androgen-intact SETs (*n* = 21); castration + ARPI SETs (*n* = 4). (**d**) The normalized gene mutation burden was calculated as the number of coding sequence single nucleotide variants (SNVs) per gene normalized for gene length per megabase (Mb)[147]. Indel burden was calculated likewise. Averaged from WES aggregated from all Stochastically Emergent Tumors (SETs) and their cognate pregrafts and visualized by chromosomal location. (**e**) Ugt1a1 gene expression difference between noncoding variant (chr1:88224769:G>T) carriers (n=7) and non-carriers (n=9) in androgen-intact samples. P-value (0.0115) by two-tailed Mann-Whitney test.

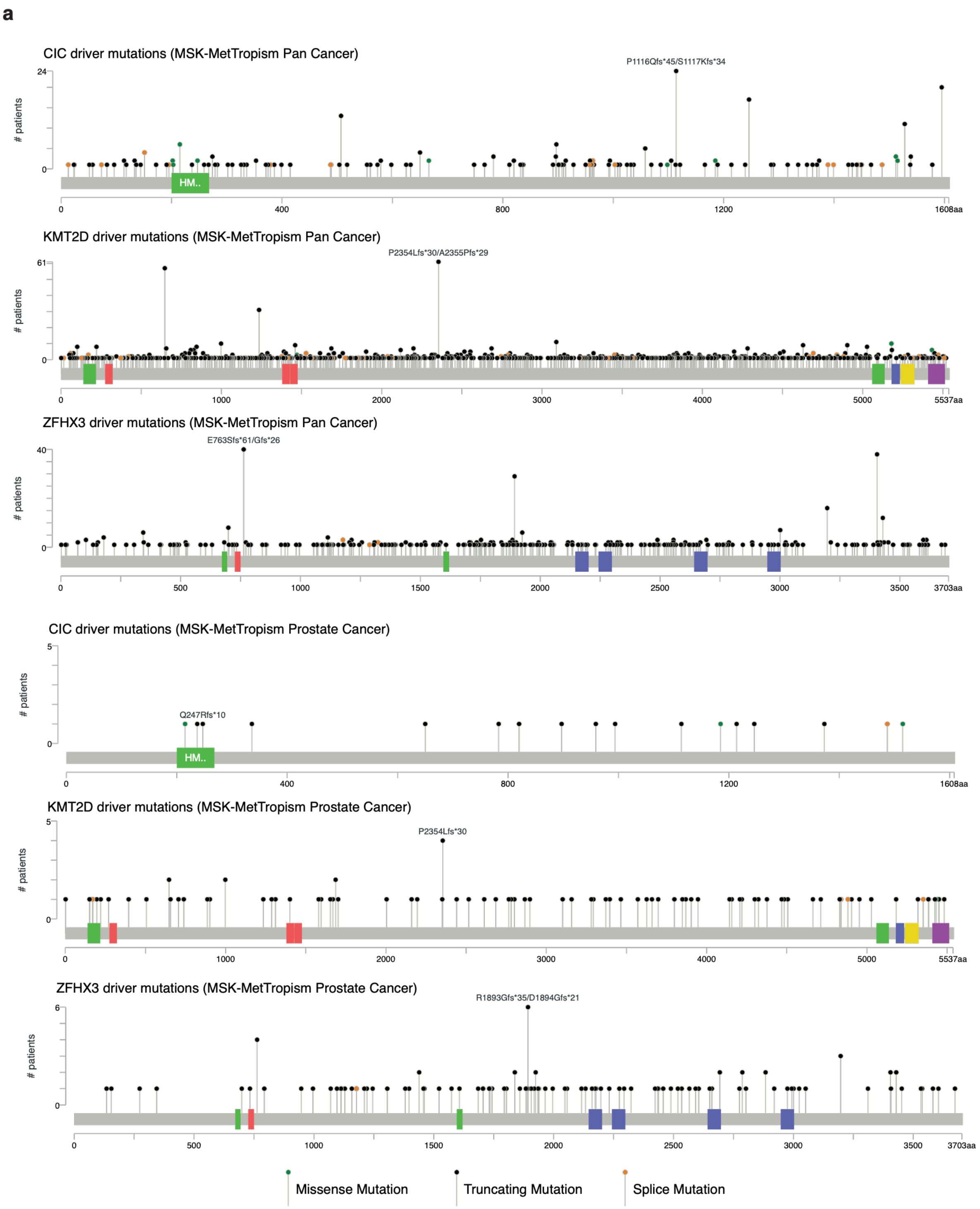


**Supplementary Figure 7 – Additional analyses of SET-identified determinants of in vivo hormone therapy response**

(**a**) Lollipop plots of putative driver mutations (annotated by OncoKB and hotspot analysis on cBioPortal) of *CIC*, *KMT2D*, and *ZFHX3* in pan cancer MSK-MetTropism cohort[33] obtained from cBioPortal[134–136]. Missense mutations (green), truncating mutations (black), and splice mutations (orange) are shown.

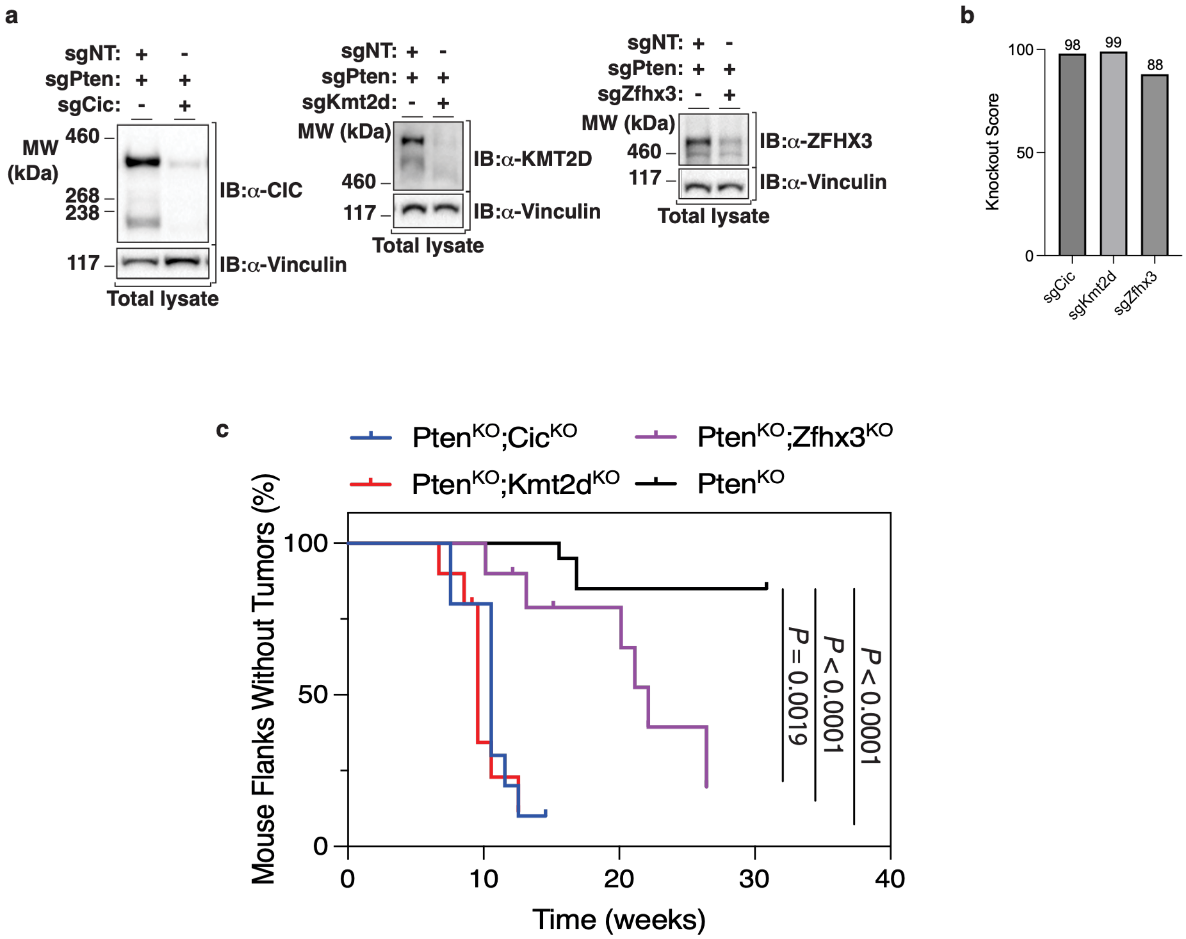


**Supplementary Figure 8 – In vivo validation of SET-identified determinants of in vivo hormone therapy response**

(**a**) Evaluation of protein levels in prostate organoid lines subjected to CRISPR knockout of putative drivers of drug response. Western blots of (*left*) CIC, (*center*) KMT2D, and (*right*) ZFHX3. Vinculin was probed as loading control. (**b**) Bar graph depicting gene knockout scores in *Pten*$^{KO}$*;Cic*$^{KO}$ (sgPten+sgCic), *Pten*$^{KO}$*;Kmt2d*$^{KO}$ (sgPten+sgKmt2d), and *Pten*$^{KO}$*;Zfhx3*$^{KO}$ (sgPten+sgZfhx3) cells when compared to *Pten*$^{KO}$ (sgPten+sgNT) control lines. Knockout scores represent the proportion of cells with either a frameshift or a 21+ base pair indel. (**c**) Tumorigenesis assay interrogating putative tumor suppressor genotypes. Kaplan-Meier curve of mouse flanks without tumors (no tumor ≥ 200 $mm^3$) following injection of prostate organoids with the indicated CRISPR knockout genotypes. P-values by Log-rank test with Holm-Šidák multiple test correction.

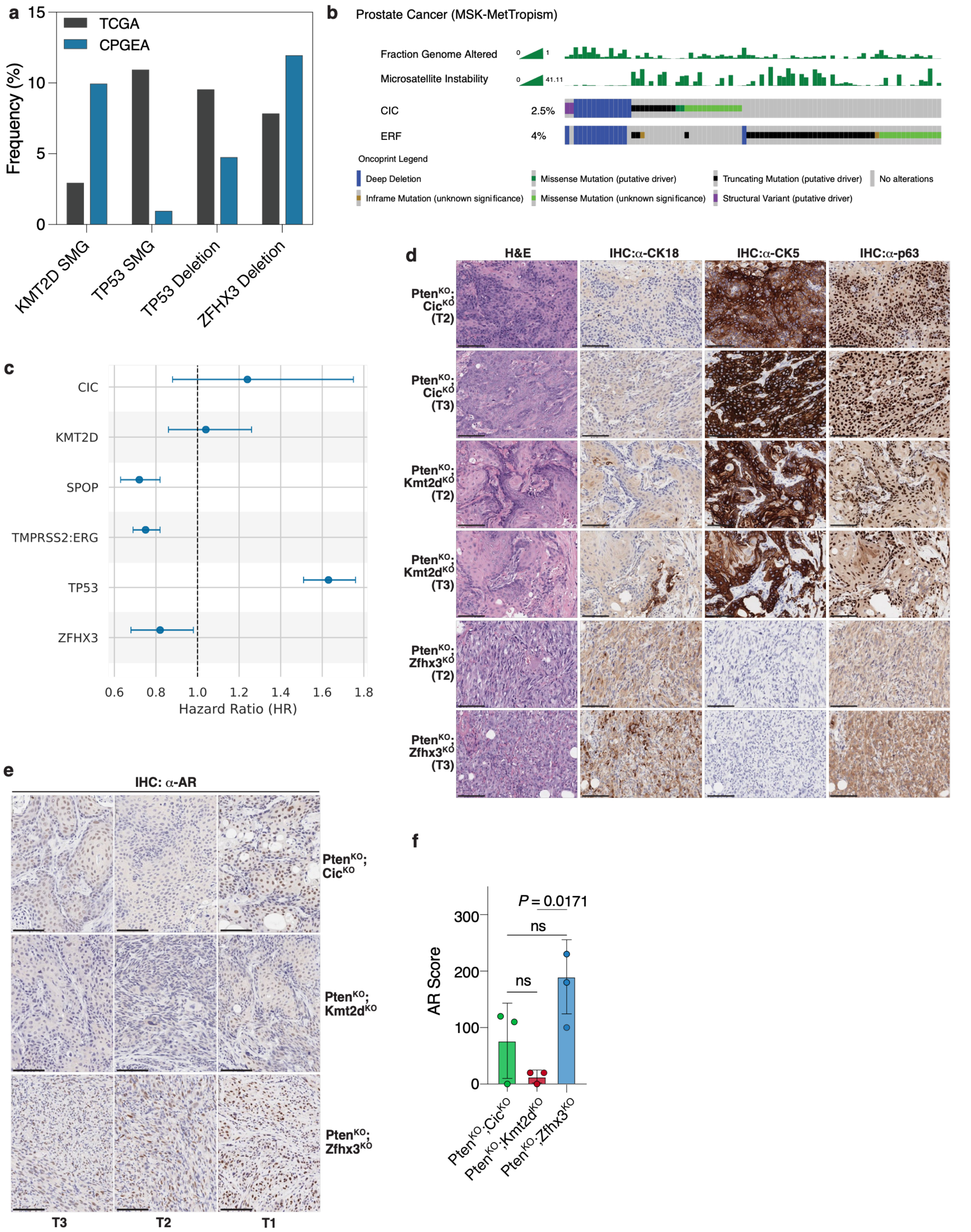


**Supplementary Figure 9 – Additional analyses of SET-identified lineage-associated hormone-response states in prostate cancer**

(**a**) Comparison of gene variant frequency between primary prostate cancer patients who are Chinese (CPGEA cohort)[148] versus those in a predominantly European American cohort (TCGA)[128]. SMG: significantly mutated gene for point mutations, per source data terminology[148]. (**b**) OncoPrint of prostate cancer patients with high FGA tumors mostly on the left and MSI-high tumors mostly on the right.  A pattern of MSI-High tumors with individually mutated *CIC* or *ERF* at an otherwise co-deleted locus is revealed. MSIsensor scores and FGA are indicated above the OncoPrint. OncoPrint is derived from prostate cancer patients in the MSK-MetTropism cohort. (**c**) Multivariate Analysis of patients with microsatellite-stable prostate adenocarcinoma profiled by Caris stratified by the presence (Group 1) or absence (Group 2) of a truncating mutation in *ZFHX3*. Covariates are TP53, CIC, KMT2D, and SPOP mutation, and TMPRSS2:ERG fusion. Hazard Ratio with 95% CI. (**d**) Hematoxylin and eosin (H&E) staining and immunohistochemistry (IHC) of androgen-intact tumors (in addition to **Figure 4d**) arising from murine injection of *Pten*$^{KO}$*;Cic*$^{KO}$, *Pten*$^{KO}$*;Kmt2d*$^{KO}$, or *Pten*$^{KO}$*;Zfhx3*$^{KO}$ prostate organoid lines. IHC antibodies target the luminal epithelial marker cytokeratin 18 (CK18) and the basal epithelial markers cytokeratin 5 (CK5) and p63. *n* = 2 distinct tumors from 2 distinct mice per genotype (Tumor 2: T2, Tumor 3: T3). Scale bars equal 100 mm. (**e**) Nuclear Androgen Receptor (AR) assessment of tumors with differential response to hormone therapy. AR IHC of representative tumors injected with *Pten*$^{KO}$*;Cic*$^{KO}$, *Pten*$^{KO}$*;Kmt2d*$^{KO}$, or sPten*;Zfhx3*$^{KO}$ organoids. Scale bars equal 100 mm. *n* = 3 distinct tumors from 3 distinct mice per genotype. (**f**) Pathologist quantification of nuclear AR among *Pten*$^{KO}$*;Cic*$^{KO}$, *Pten*$^{KO}$*;Kmt2d*$^{KO}$, or *Pten*$^{KO}$*;Zfhx3*$^{KO}$ tumors. A genitourinary pathologist assessed an AR score for each tumor based on the nuclear AR intensity (0-3+) multiplied by the percentage of AR positive cells. P-value by one-way ANOVA with Tukey post hoc test (ns, not significant).
